# Actin cytoskeleton signaling via MRTF/SRF entrains circadian clock

**DOI:** 10.1101/2022.04.29.490111

**Authors:** Xuekai Xiong, Weini Li, Jin Nam, Meng Qu, Steve A. Kay, Ke Ma

## Abstract

The circadian clock is entrained to daily environmental cues. Integrin-linked intracellular signaling *via* actin cytoskeleton dynamics transduces extracellular matrix interactions to Myocardin-related Transcription Factor (MRTF)/Serum Response Factor (SRF)-mediated transcription. Actin cytoskeleton organization in liver displays diurnal oscillations and SRF-MRTF activity exert transcriptional control to entrain clock. By interrogating disparate upstream events involved in actin cytoskeleton-MRTF-A/SRF signaling cascade, here we show that this signaling cascade transduce cellular niche cues to modulate circadian clock function. Pharmacological inhibitions of MRTF-A/SRF, by disrupting actin polymerization or blocking ROCK kinase, induced period lengthening with augmented clock amplitude, and genetic loss-of-functions of *Srf* or *Mrtf-a* mimic that of actin-depolymerizing agents. In contrast, actin-polymerization induced by Jasplakinolide shortened period with attenuated amplitude. In addition, interfering with cell-matrix interaction through blockade of integrin, inhibition of focal adhesion kinase or attenuating matrix rigidity led to reduced period length while enhancing amplitude. Mechanistically, we identify that core clock repressors, Per2, Nr1d1, and Nfil3, are direct transcriptional targets of MRTF-A/SRF in mediating actin dynamic-induced clock response. Collectively, our findings defined an integrin-actin cytoskeleton-MRTF/SRF pathway in linking clock entrainment with extracellular microenvironment that may facilitate cellular adaptation to its physical niche.

**Summary statement:** Our study revealed the role of actin cytoskeleton-MRTF/SRF signaling in entraining circadian clock to its extracellular physical niche environment.

## Introduction

The circadian clock has evolved as a time-keeping mechanism to anticipate and adapt to daily environmental changes essential for organismal fitness and survival (Finger et al., 2020; Takahashi, 2017). Under physiological conditions, a central clock pacemaker residing in suprachiasmatic nuclei (SCN) of the hypothalamus is entrained by daily light input that synchronizes clock circuits within peripheral tissues. In additional to SCN-driven signals for synchronization, cell-autonomous peripheral oscillators in tissues outside SCN respond to humoral signals and distinct tissue-specific cues (Dibner et al., 2010; Finger et al., 2020; Schibler and Sassone-Corsi, 2002). Synchronization of clock circuits in the body ensures coordinated temporal orchestration of daily physiological processes. Disruption of clock synchrony, such as shiftwork or jet-lag, predispose to various disease risks including the development of cancer (Kettner et al., 2014; Lin and Farkas, 2018; Mocellin et al., 2018) and metabolic disorders (Buxton et al., 2012; Karatsoreos et al., 2011; Scheer et al., 2009; Shi et al., 2013; Turek et al., 2005). Better understanding of the external or endogenous stimuli that entrains the circadian clock provides fundamental knowledge of the mechanisms that underly environmental adaptation of the circadian clock (Finger et al., 2020).

The molecular circuit driving the ∼24-hour rhythm of circadian oscillators is composed of a transcription-translation feedback loop (Takahashi, 2017). Circadian Locomotor Output Kaput (CLOCK) and Brain and muscle Arnt-like 1 (Bmal1), the key transcriptional activators of this molecular clock feedback loop, heterodimerize and activate the transcription of clock repressor proteins, the Periods (Per1, Per2 & Per3) and Cryptochromes (Cry1 and Cry2). Ensuing cytosolic accumulation, phosphorylation and nuclear translocation of the Period and Cryptochrome proteins inhibit Bmal1/CLOCK-dependent transcription *via* direct interaction with the heterodimer. This negative transcriptional feedback cycle coupled with translational control constitutes the core clock regulatory loop. An additional Rev-erbα/ROR-mediated Bmal1 transcriptional oscillation re-enforces the robustness of this core clock mechanism (Preitner et al., 2002).

Daily entrainment to environmental cues is essential for internal synchronization of the body clock system and adaptation to cyclic changes (Finger et al., 2020). Despite our current knowledge of the intricate molecular network driving clock oscillation and entrainment, how clock respond and adapt to its immediate niche environment is yet to be addressed. Various extracellular physical or chemical cues, including cell adhesion to extracellular matrix (ECM) *via* integrin or growth factor activation *via* cell-surface receptors, are transduced intracellularly by a signaling cascade that involves actin cytoskeleton remodeling and downstream transcriptional response mediated by serum response factor (SRF) in response to Myocardin-related Transcription Factor (MRTF) activation (Olson and Nordheim, 2010; Posern and Treisman, 2006). Integrins, through direct interactions with specific extracellular matrix (ECM) components, form focal adhesion complexes that connect the cellular physical environment with intracellular actin cytoskeleton network through a myriad of signaling pathways (Romero et al., 2020). Activation of integrin-mediated intracellular signaling transduction, including focal adhesion-associated kinase (FAK), Rho-GTPases, ROCK kinase and their associated effector molecules, transmits extracellular microenvironment cues to modulate actin polymerization *via* the formation of filamentous actin (F-actin) from monomeric globular actin (G-actin). Various growth factors or cytokines, including PDGF, TGF-β or Wnt, also elicit Rho-GTPase/ROCK signaling through cell surface receptors to engage actin cytoskeleton dynamic regulation. Actin polymerization leads to the release of Myocardin-related Transcription Factors (MRTF-A/B, MKL-1/2) from sequestration by G-actin monomers, with subsequent nuclear translocation and activation of SRF-mediated transcription (Posern and Treisman, 2006). Nuclear shuttling of MRTF, in response to actin dynamic regulation, interacts with SRF on its cognate DNA binding motif, the CArG box, to control target gene expression (Miano, 2003). MRTF/SRF-controlled genes participate in various physiological processes during tissue development, growth and remodeling involving cytoskeleton organization, such as matrix adhesion, migration, proliferation and differentiation (Gualdrini et al., 2016; Long et al., 2007; Olson and Nordheim, 2010; Wang et al., 2002).

Serum is known as an universal synchronizing signal of cellular clocks (Balsalobre et al., 1998). Serum-stimulated intracellular actin turnover entrains the liver clock (Gerber et al., 2013), while MRTF/SRF mediates the transcriptional response to actin dynamic-elicited clock entrainment (Esnault et al., 2014). Despite the role of intracellular actin dynamic and MRTF/SRF activity in mediating cellular interactions with ECM, it remains unclear whether extracellular niche impacts circadian clock function. In the current study, we employed pharmacological and genetic approaches targeting disparate steps of integrin-actin cytoskeleton-MRTF/SRF signaling transduction to establish its role in modulating clock oscillation.

## Results

### Pharmacological perturbations of actin polymerization modulate clock oscillation

Polymerization of monomeric actin, in response to upstream signals, is the key regulatory step that releases MRTF from G-actin sequester to activate SRF-mediated transcription (Olson and Nordheim, 2010). To test whether actin dynamic-associated signaling modulates circadian clock, we first determined the effects of pharmacological agents that disrupt or promote actin polymerization using mouse fibroblasts containing a Period-2 promoter-driven luciferase reporter knock-in (Per2::Luc) (Yoo et al., 2004). Continuous monitoring of bioluminescence activity of Per2::Luc fibroblasts revealed that Cytochalasin D (Cyt D), a known actin depolymerization compound (Hill et al., 1995; Wei et al., 2001), significantly altered clock cycling properties, as shown by average tracing for 5 days (Fig. 1A & 1B). Cyt D, within 1-5μM tested, induced longer period length (Fig. 1C) and a dose-dependent increase in oscillation amplitude (Fig. 1D). The effect of Cyt D on actin organization was demonstrated using phalloidin immunofluorescence staining of F-actin stress fibers. Cty D-treated fibroblasts displayed marked reductions of F-actin that persisted at 24 hours after treatment, with cells viable at both concentrations tested (Fig. 1E). The loss of actin polymerization is consistent with round up cell morphology observed (Suppl. Fig. S1). As expected of Cyt D action in interfering with actin polymerization and consequent inhibition of MRTF/SRF activity, expression of known MRTF/SRF target genes, connective tissue growth factor (*Ctgf*) and Four and half LIM-domain protein 1 (*Fhl1*), were markedly down-regulated by Cyt D (Fig. 1F). *Mkl1* transcript was also reduced, although *Srf* expression was not altered. Analysis of core clock genes revealed significant effects of Cyt D on inhibiting *CLOCK, Per2* and *Cry2* expression, without altering other components (Fig. 1G). Down-regulation of clock genes by Cyt D suggests potential transcriptional regulation *via* actin cytoskeleton-related signaling, and dampening of positive clock component with augmented negative regulatory loop may contribute to the period lengthening effect of Cyt D. Using U2OS cells with Per2::Luc reporter, we further examined CytD effect on clock modulation and found it induced similar dose-dependent period-lengthening (Fig. 1H & 1I), although with reduced cycling amplitude at 5μM (Fig. 1J).

**Figure 1.**
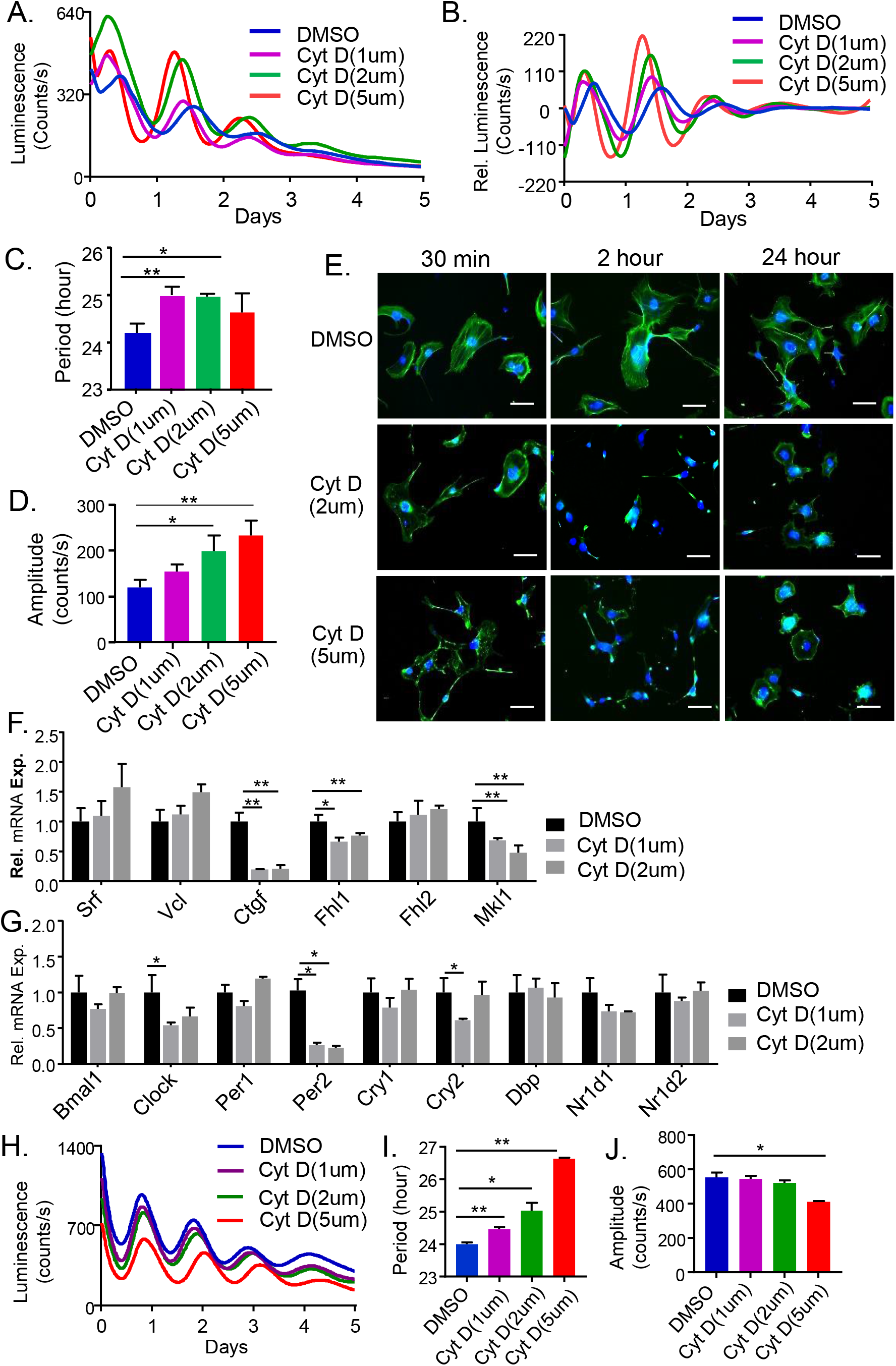
Effect of Cytochalasin D on clock rhythm. (A, B). Bioluminescence recordings of average original (A), and baseline-subtracted (B) plots of Per2-Luciferase knock-in reporter in mouse fibroblasts subjected to Cytochalasin D (Cyt D) treatment at indicated concentrations, with quantitative analysis of clock period length (C) and amplitude (D). N=3/ treatment group. (E) Representative fluorescence images of F-actin stress fibers by phalloidin staining in control (DMSO) and Cyt D-treated fibroblasts. Scale bar: 100μm. (F & G) RT-qPCR analysis of Cyt D effect on expression of target genes and signaling components of MRTF/SRF-mediated transcription (F), and core clock genes (G, n=3). (H-J) Effect of Cytochalasin D on clock modulation in Per2::Luc U2OS cells. Bioluminescence recording of the average original plots of Per2-Luciferase knock-in U2OS cells subjected to Cytochalasin D at indicated concentrations (H), with quantitative analysis of clock period length (I) and amplitude (J). N=3/treatment group. *, **: P≤0.05 or 0.01 Cyt D vs. DMSO by Student’s t test.

We next determined whether additional molecules that inhibit actin polymerization, such as Latrunculin B (Lat B), display shared clock-modulatory activity with Cyt D (Allingham et al., 2006). Indeed, Lat B led to significant period lengthening at 2 uM (Fig. 2A-2C), with dose-dependent effect on increasing amplitude (Fig. 2D). Lat B appear to induce phase advance as compared to Cyt D (Fig. 2B), although this was not directly assessed with re-synchronization. The loss of F-actin organization induced by Lat B was more rapid and robust than that of Cyt D, with nearly abolished F-actin staining of at 30 minutes upon treatment and persisted at 24 hours (Fig. 2F). Lat B effect on actin cytoskeleton is also reflected in the rapid change of cell morphology (Suppl. Fig. S1). Lat B resulted in marked inhibition of SRF target genes at concentrations tested, with near complete inhibition of *Ctgf* and strong suppression of *Srf, vinculin* (*Vcl*), *Fhl1* and *Fhl2* (Fig. 2F). Lat B led to significant down-regulation of *Bmal1*, while *Per1, Dbp* and *Nr1d2* were induced at 5 μM concentration (Fig. 2G). In U2OS cells, Lat B induced comparable effects on increasing period and amplitude (Fig. 2H-2I). Additionally, we tested another compound that disrupts actin cytoskeleton, Cytochalasin B, and found comparable effects in prolonging period with augmentation of cycling amplitude in Per2::Luc mouse fibroblasts (Fig. S2).

**Figure 2.**
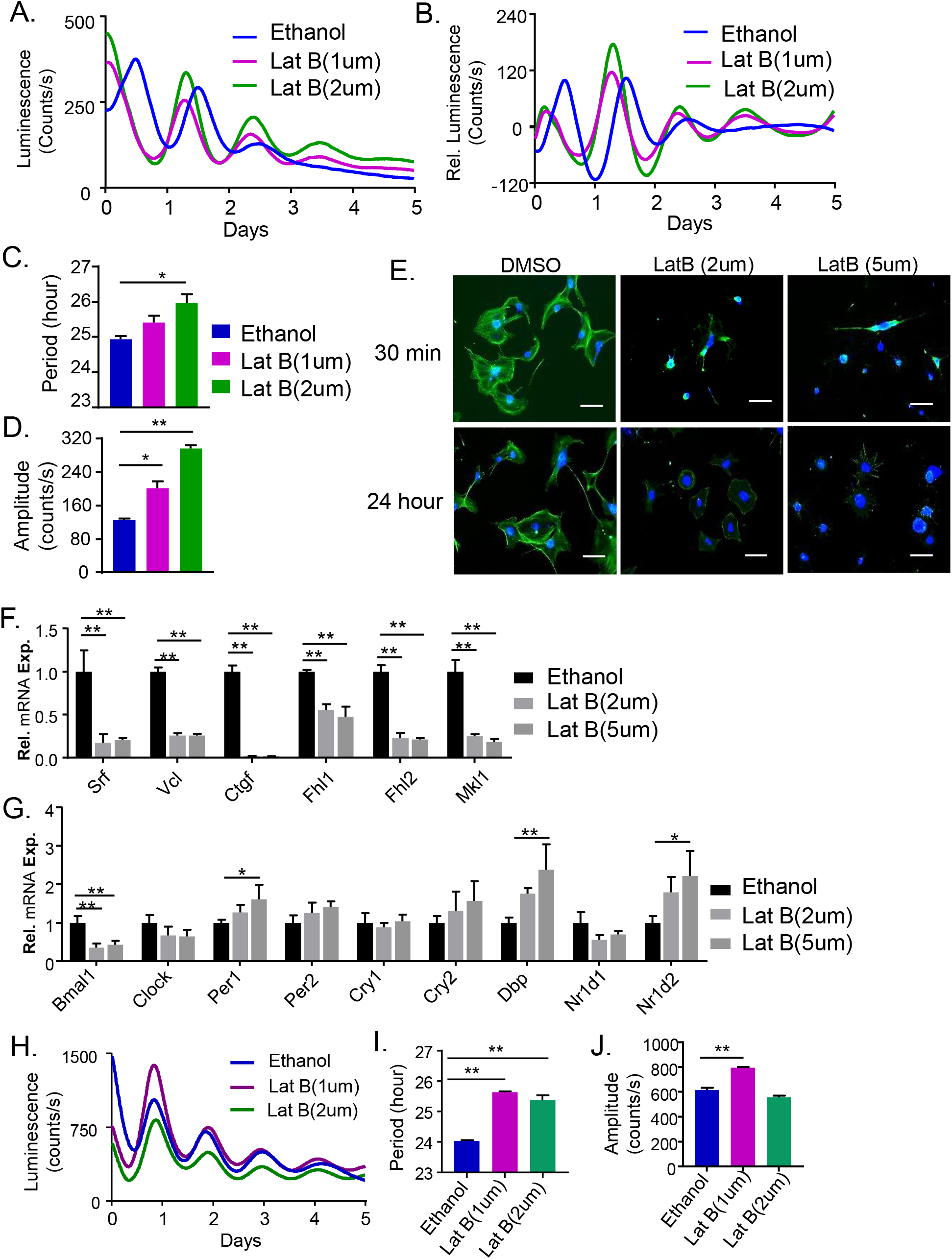
Effect of Latrunculin B on clock rhythm modulation. (A-D) Bioluminescence recordings of the average of original (A), and baseline-subtracted plots (B) of Per2-Luciferase knock-in fibroblasts subjected to Latrunculin B (Lat B) treatment at indicated concentrations, with quantitative analysis of clock period length (C) and amplitude (D, n=3). (E) Representative fluorescence images of phalloidin staining of F-actin in control (DMSO) and Lat B-fibroblasts. Scale bar: 100μm. (F, G) RT-qPCR analysis of the effect of Lat B on expression of MRTF/SRF-related signaling pathways (F), and core clock genes (G, n=3). (H-I) Effect of Latrunculin B on clock in Per2::Luc U2OS cells. Bioluminescence recording of the average original plots of Per2-Luciferase U2OS cells subjected to Lat B at indicated concentrations (H), with quantitative analysis of clock period length (I) and amplitude (J). N=3/treatment group. N=3/treatment group. *, **: P≤0.05 or 0.01 Lat B vs. DMSO by Student’s t test.

Furthermore, we tested whether promoting actin polymerization *via* Jasplakinolide (Jas) (Allingham et al., 2006), thereby stimulating MRTF/SRF-mediated transcription, affects clock function. In contrast to Cyt D, Jas resulted in dose-dependent period shortening from 0.1 to 0.5 μM (Fig. 3A-C), with significantly decreased oscillation amplitude at 0.5 μM (Fig. 3D). Despite limited effect of Jas on augmenting F-actin (Fig. 3E), potentially due to robust F-actin in fibroblasts under normal culture conditions, gene expression analysis revealed activation of SRF-mediated transcription, as shown by marked inductions of *Vcl, Ctgf, Fhl1, Fhl2* and *Srf* itself (Fig. 3F). As compared to the suppression of clock repressors by Cyt D, Jas elicited up-regulations of *Bmal1, Per2, Cry2, Nr1d1* and *Nr1d2* in a dose-dependent manner (Fig. 3G), although *Per1* was reduced. Notably, Jas had stronger effect on reducing period length and clock amplitude in U2OS cells containing Per2::Luc starting at 0.5 μM (Fig. 3H-3I). Thus, promoting actin polymerization exert opposing effects on clock cycling properties as compared to actin-depolymerization.

**Figure 3.**
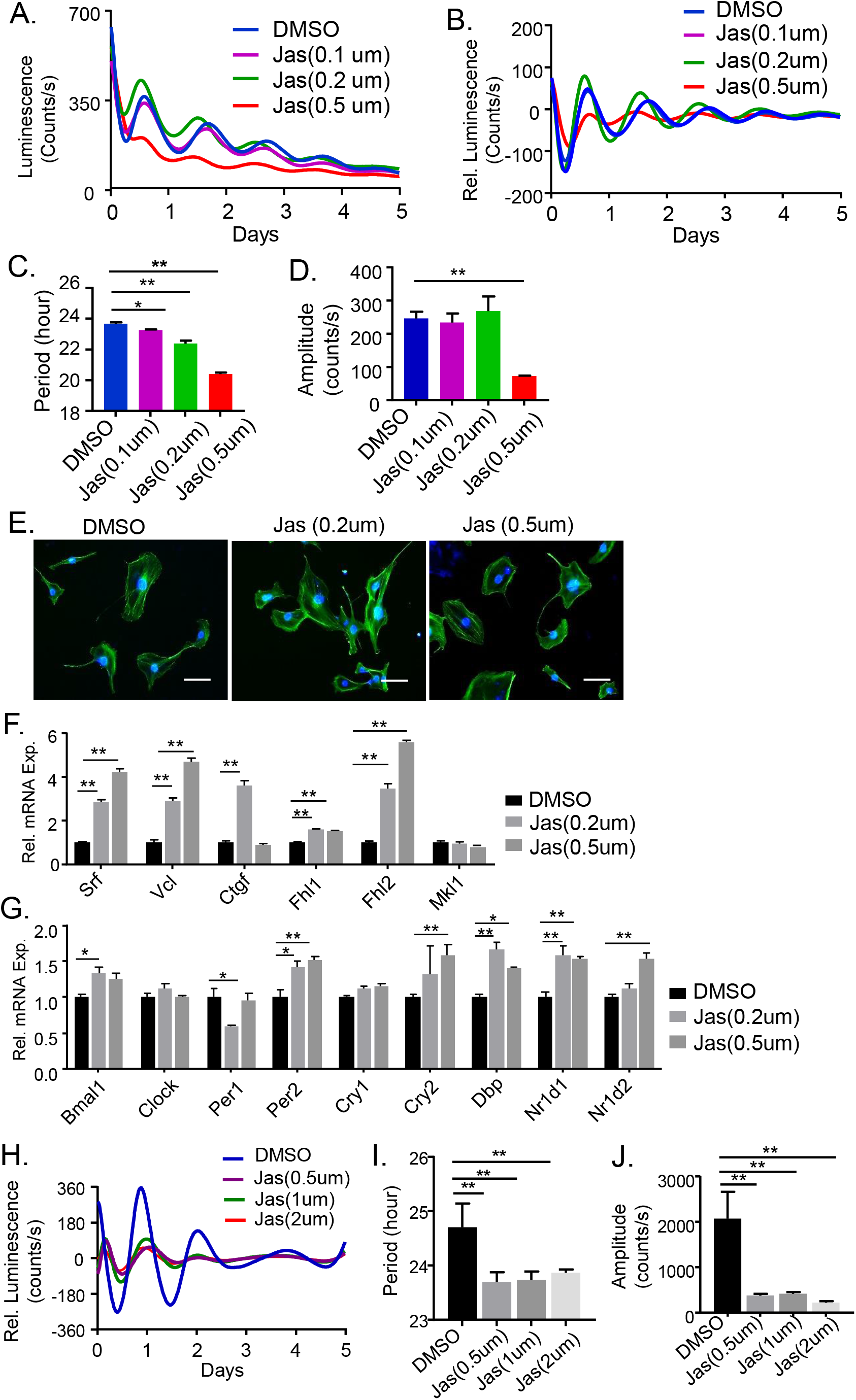
Effect of actin-polymerizing Jasplakinolide on clock modulation. (A-D) Average original bioluminescence recordings (A) and baseline-subtracted plots (B) of Per2-Luciferase activity in mouse fibroblasts subjected to Jasplakinolide (Jas) treatment at indicated concentrations, with quantitative analysis of clock period length (C) and amplitude (D). N=3/treatment group. (E) Representative fluorescence images of phalloidin staining (green) of F-actin in control (DMSO) and Jas-treated fibroblasts for 2 hours. Scale bar: 100μm. (F & G) RT-qPCR analysis of the effect of Jas on expression of MRTF/SRF signaling components (F), and core clock genes (G). N=3. (H-I) Effect of Jasplakinolide on clock in Per2::Luc U2OS cells. Bioluminescence recording of the average baseline-subtracted plots of Per2-Luciferase U2OS cells subjected to Jas at indicated concentrations (H), with quantitative analysis of clock period length (I) and amplitude (J). N=3/treatment group. N=3/treatment group. *, **: P≤0.05 or 0.01 Jas vs. DMSO by Student’s t test.

### Genetic inhibition of SRF or MRTF function modulates clock property

Intracellular signaling induced by actin dynamics ultimately results in MRTF/SRF-mediated transcription activation response. Using siRNA-mediated silencing of these effectors of actin dynamics, we tested whether SRF or MRTF-A mediates the effects of actin cytoskeleton-modifying molecules on clock modulation. One among three *Srf* siRNA tested was effective in abolishing SRF protein expression (Fig. 4A). *Srf* silencing altered clock oscillatory properties similarly as actin-depolymerizing agents (Fig. 4B), with increased period length (Fig. 4C) and tendency toward higher amplitude (Fig. 4D). We synchronized the cells by serum to determine effect of SRF inhibition on clock genes at two circadian times, CT12 (CT: circadian time) and CT24. *Mkl1, Fhl1* and *Fhl2* displayed circadian time-dependent regulations with higher expressions at CT24 than that of CT12 (Fig. 4E). Knockdown of *Srf* markedly reduced *Ctgf* and *Fhl1* expression at both time points, as compared to that of the scrambled controls (SC), while *Mkl1* and *Fhl2* levels were significantly lower only at CT24. Analysis of clock genes revealed inductions of *Per1* and *Cry1* at CT24 than that of CT12, while both were significantly down-regulated by *siSrf* at CT24 (Fig. 4F). Furthermore, loss of *MRTF-A* by siRNA knockdown (*siMrtf*) resulted in similar effects as SRF inhibition. Two *siMrtf* examined largely abolished MRTF-A protein expression (Fig. 4G), and both resulted in increased period length (Fig. 4H & 4I) with augmented amplitude (Fig. 4J) similar to *Srf* inhibition. These findings indicate that SRF and MRTF are involved in clock function with transcriptional regulation of the core clock circuit.

**Figure 4.**
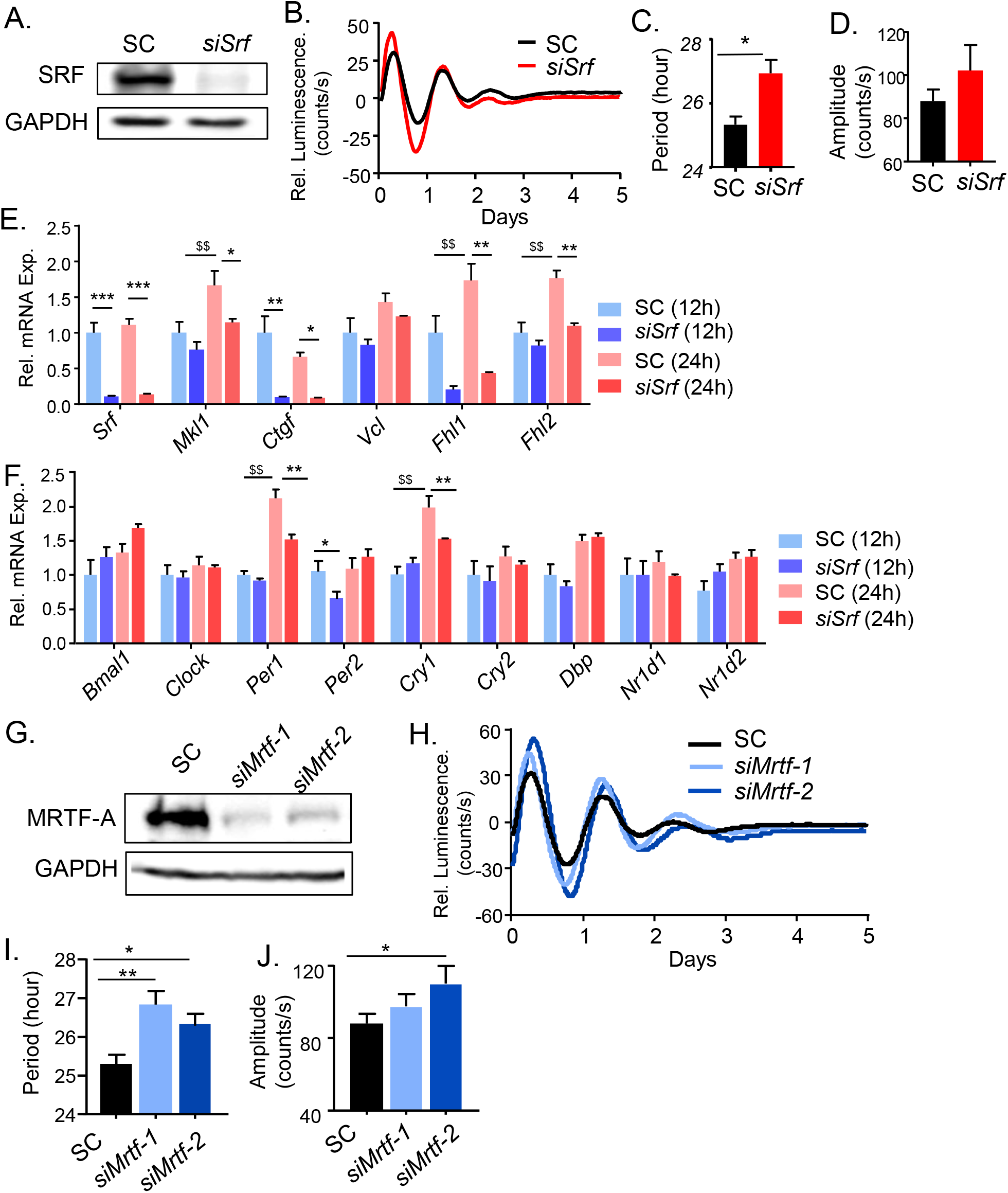
Genetic loss-of-functions of SRF and MRTF-A influence clock function. (A-F). Effect of *Srf* silencing on core clock properties. (A) Immunoblot analysis of SRF protein expression in Per2-Luc fibroblasts with transient knockdown of scrambled control (siSC) or *siSrf*. (B-D) Representative bioluminescence tracings of Per2-Luc fibroblasts (B) with quantification of clock period length (C) and amplitude (D). N=3. *, **: P≤0.05 or 0.01 *siSrf* vs. SC. (E & F) RT-qPCR analysis of MRTF/SRF-related signaling components (E), and core clock gene expression (F) at 12 and 24 hours after serum shock synchronization (n=3). $, $$: P≤ 0.05 or 0.01 control 24hr vs. 12hr; and *, **: P≤0.05 or 0.01 *siSrf* vs. SC. (G-J) Effect of MRTF-A silencing on clock activity. Immunoblot analysis of MRTF-A protein level (G). (H) Representative bioluminescence of Per2-Luc fibroblasts transfected with scrambled control (SC) or *siMrtf*, with quantification of clock period (I) and amplitude (I). N=3. *, **: P≤0.05 or 0.01 *siMrtf* vs. SC by student’s t test.

### Inhibition of upstream signaling pathways inducing actin cytoskeleton remodeling modulates clock function

The Rho GTPases (RhoA, Rac1 and Cdc42) and Rho-associated kinase (ROCK) are upstream signaling molecules transmitting cell surface stimuli to modulate actin dynamics (Hill et al., 1995; Wei et al., 2001). Cell surface ECM or biochemical cues, *via* integrin or receptor tyrosine kinases, activate Rho GEFs and Rho/ROCK-mediated kinase cascade to alter G-actin to F-actin ratio (Esnault et al., 2014). We thus tested whether perturbing Rho GTPase signaling upstream of actin dynamics by a specific ROCK kinase inhibitor, Y27632, impacts clock function. As shown in Fig. 5A-5D, Y27632 exerted similar effects as actin-disrupting molecules, with dose-dependent lengthening of period (Fig. 5C) accompanied by higher amplitude (Fig. 5D). Y27632 induced a robust but delayed effect on F-actin as compared to actin depolymerizing agents, as shown by cell shape change and attenuated phalloidin staining most evident at 2 hours following treatment (Fig. S3). As expected with inhibition of MRTF/SRF activity, Y27632 markedly down-regulated SRF targets with nearly ∼80% lower mRNA expressions of *Srf, Vcl, Ctgf* and *Mkl1*, as compared to DMSO-treated controls (Fig. 5E). Y27632 also suppressed *Per2, Nr1d1* and *Nr1d2* expression similar to that of Cyt D (Fig. 5F), whereas its induction of *Cry1* and *Cry2* are distinct. Thus, perturbing Rho GTPase signaling up-stream of actin remodeling largely recapitulated the effects of actin depolymerization on clock modulation. In comparison, Y27632 effect on period length was moderate in U2OS cells, with significantly prolonged period and increased amplitude only observed at 50uM (Suppl. Fig. S4).

**Figure 5.**
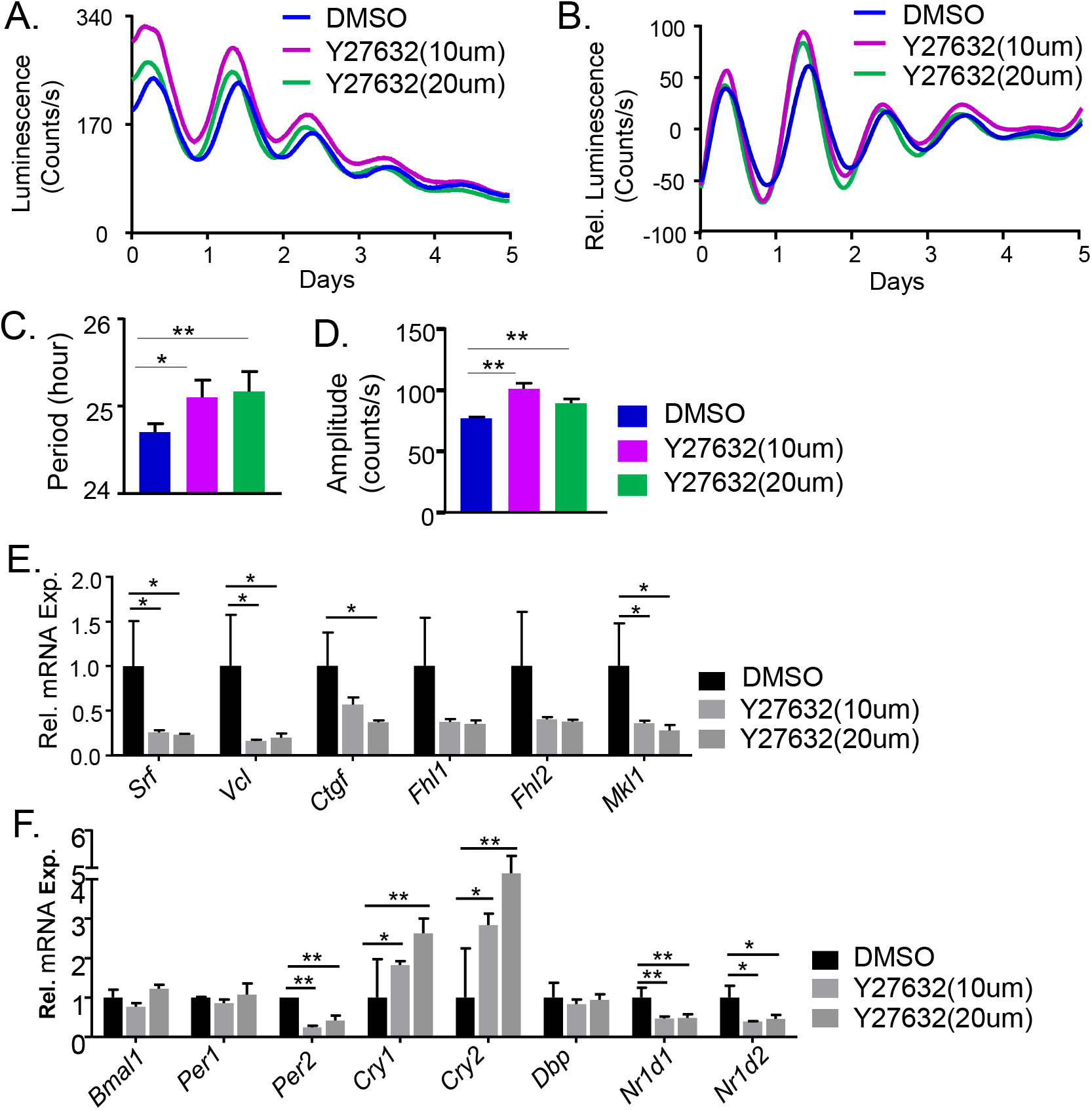
Effect of ROCK inhibition by Y27632 and MLCK inhibition by ML7 on circadian clock properties. (A-E). Representative raw bioluminescence (A), and baseline-subtracted plots (B) of dose-dependent effects of ROCK inhibitor Y27632 on Per2-Luciferase activity, with quantitative analysis of clock period length (C) and cycling amplitude (D). N=3. (E & F) RT-qPCR analysis of the effect of Y27632 on MRTF/SRF signaling pathway (E), and core clock genes (F, n=3). *, **: P≤0.05 or 0.01 Y27632 vs. DMSO control by student’s t test.

### Integrin-mediated adhesion signaling and ECM stiffness modulate clock function

ECM constitutes the immediate physical micro-environment cells residing in, and integrin-mediated focal adhesion complex links ECM components with intracellular actin cytoskeleton (Olson and Nordheim, 2010). Focal adhesion kinase (FAK), together with additional components of the focal adhesion complex, transduces ECM cues to modulate F-actin stress fiber formation with activation of MRTF-SRF transcriptional response (Romero et al., 2020). Based on the finding that actin dynamic-induced MRTF/SRF activity modulates circadian clock, we postulated that blocking integrin-ECM interaction and associated focal adhesion signaling may impact clock function and thus tested the effect of integrin blockade via integrin αV-targeting cyclic peptide (cRGD). As compared to a peptide control with scrambled sequence, cRGD-treated cells displayed a tendency toward longer period (Fig. 6A & 6B) with significantly higher oscillation amplitude at 2uM (Fig. 6C). Down-regulations of SRF transcriptional targets, *Srf, Vcl* and *Ctgf*, revealed inhibition of MRTF/SRF signaling by cyclo-RGD (Fig. 6D), together with suppressed *Bmal1* and *Per2* as compared to control peptide-treated cells (Fig. 6E). To further test integrin-induced intracellular signaling through FAK, we used siRNA knockdown targeting FAK (*siFAK)*, with demonstrated efficiency in attenuating FAK protein level (Fig. 6F). All three FAK silencing tested led to consistently prolonged clock period (Fig. 6G & 6H), although *siFAK* did not affect cycling amplitude (Fig. 6I).

**Figure 6.**
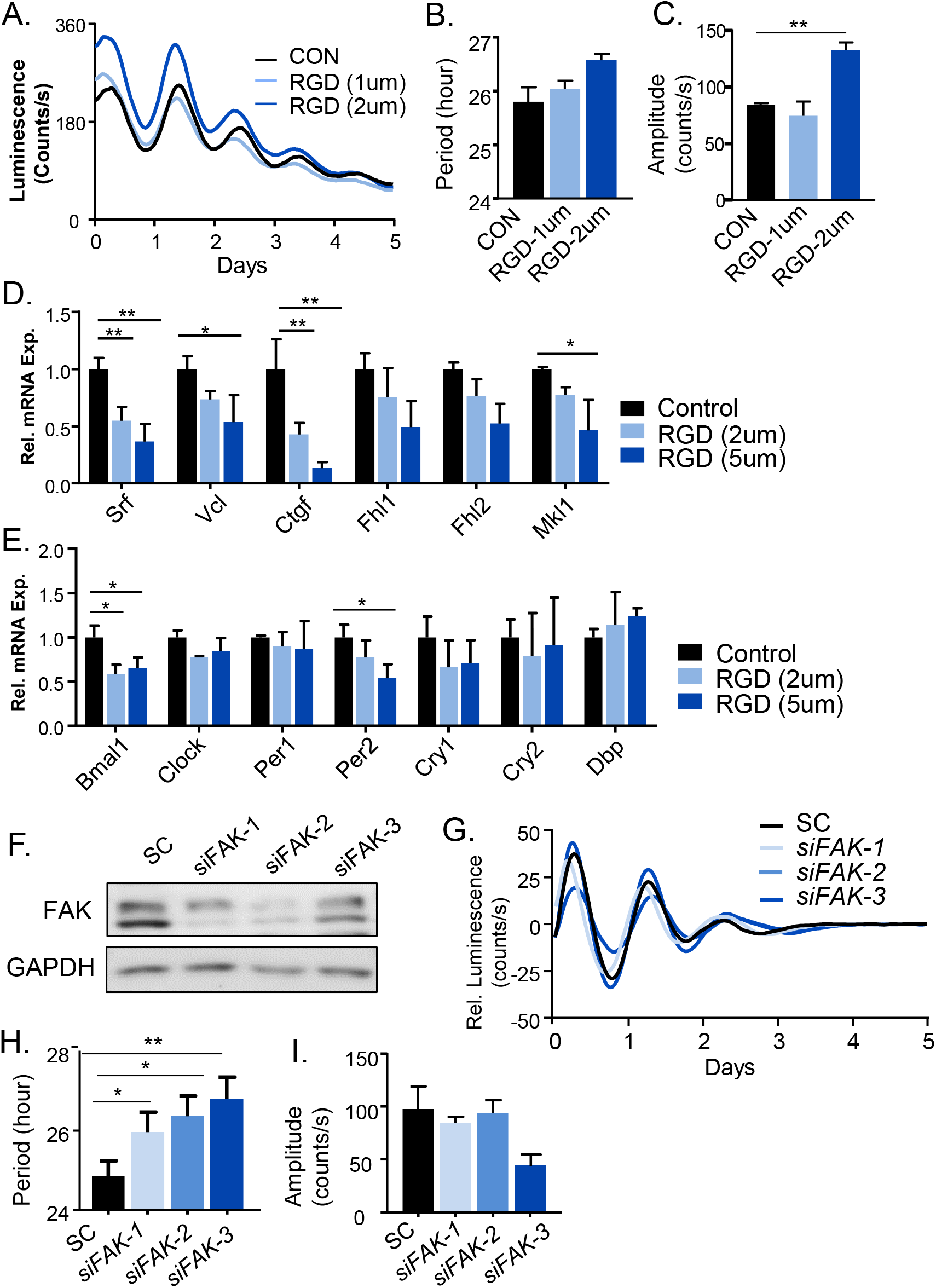
Inhibition of integrin-mediated focal adhesion signaling modulates clock function. (A-E) Effect of integrin blocking peptide Cyclo RGD on clock function. Representative bioluminescence of dose-dependent effects of RGD on Per2-Luciferase activity (A), with quantitative analysis of clock period length (B) and cycling amplitude (C, n=3). (D & E) RT-qPCR analysis of the effect of integrin blocking peptide on MRTF/SRF signaling components (D), and core clock gene expression (E, n=3). *, **: P≤0.05 or 0.01 RGD blocking peptide vs. control peptide. (F-I) Effect of focal adhesion kinase (FAK) gene silencing by siRNA knockdown on clock oscillation. Immunoblot analysis of FAK protein level (F), representative bioluminescence of Per2-Luc fibroblasts transfected with scrambled control (SC) or *siFAK* (G), with quantification of clock period length (H) and amplitude (I, n=3). *, **: P≤0.05 or 0.01 *siFAK* vs. SC by student’s t test.

To determine whether physical properties of ECM has direct impact on clock function, we applied two engineered electrospun scaffolds with differing stiffness to Per2-Luc fibroblasts culture. A soft polycaprolactone (PCL) matrix with 19 kPa elastic modulus (EM) and a stiff ∼300 kPa polyether ketone ketone (PEKK) were selected to test ECM rigidities analogous to soft adipose tissue and bone-like tissue respectively, as previously described (Maldonado et al., 2016; Maldonado et al., 2017; Maldonado et al., 2015). In comparison to the much more rigid standard polystyrene tissue culture plastic with EM >1000 kPa as control, both engineered scaffolds prolonged clock period length, with PCL inducing a longer period than PEKK (Fig. 7A & 7B). Clock cycling amplitude on these matrices were significantly augmented (Fig. 7C). Similar clock response to matrix stiffness were obtained using U2OS containing Bmal1 promoter-driven luciferase reporter, with soft PCL resulting in period lengthening and PEKK augmenting clock amplitude (Fig. 7D-7F). These effects on clock were in line with findings from actin-disrupting chemicals, suggesting that reduced ECM tension could be transduced, potentially *via* actin cytoskeleton dynamics, to influence clock activity.

**Figure 7.**
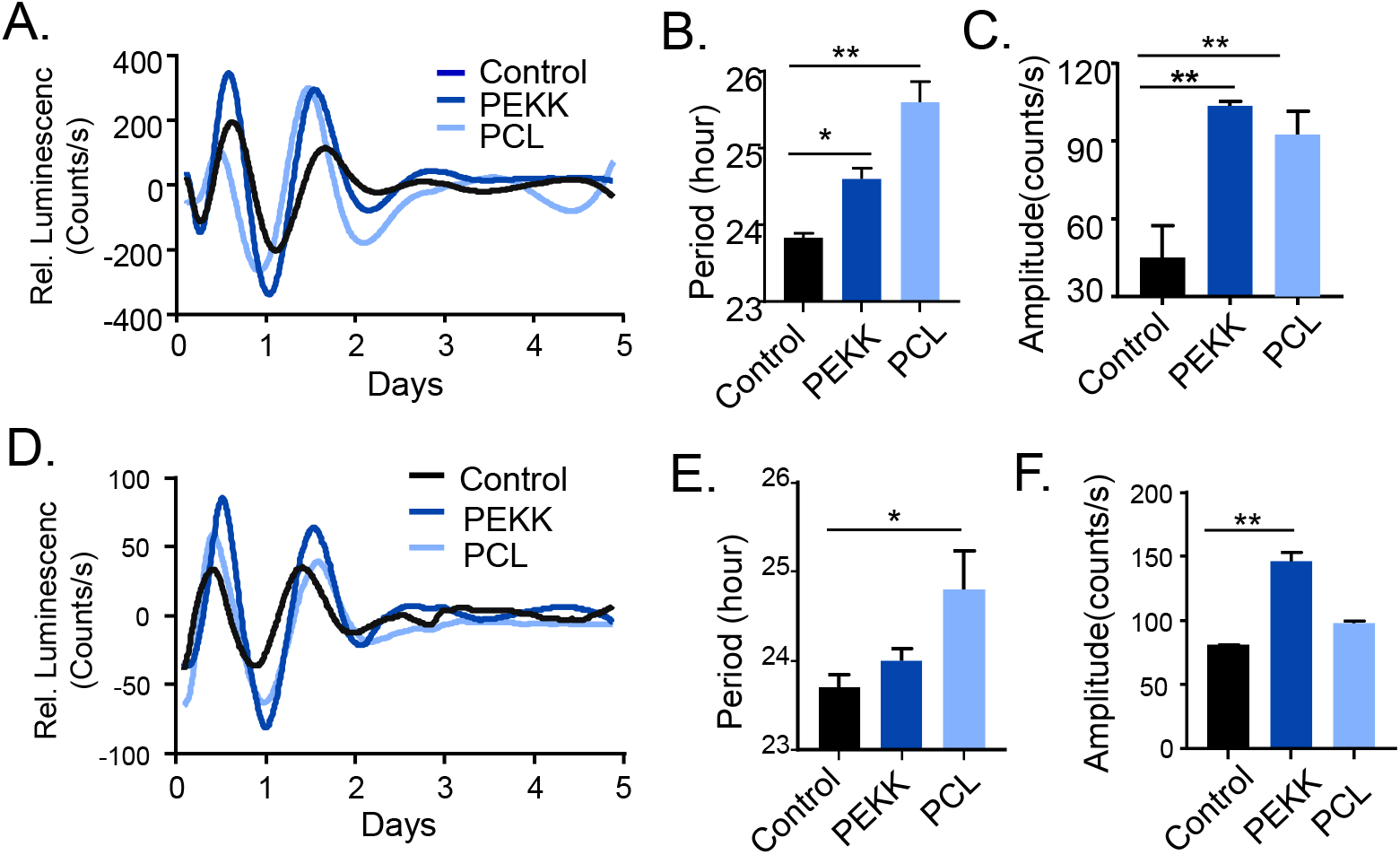
Effect of extracellular matrix stiffness on clock modulation. (A-C) Average bioluminescence baseline-subtracted tracing of Per2-Luciferase mouse fibroblasts on electrospun polyether ketone ketone (PEKK, 300 kPa elastic modulus) and polycaprolactone (PCL, 20 kPa) scaffolds with tissue culture polystyrene as control (A), with quantitative analysis of clock period length (B) and cycling amplitude (C). N=4/group. (D-F) Average bioluminescence tracing of Per2-Luc reporter in U2OS cells on PCL and PEKK scaffolds (D), with quantitative analysis of clock period length (E) and cycling amplitude (F, n=4).

### MRTF and SRF exerts direct transcriptional control of core clock regulators

In response to altered G-actin/F-actin ratio, MRTF translocates to the nucleus and binds SRF to activate gene transcription *via* a consensus CArG box motif response element (Sun et al., 2006). While SRF association with cognate DNA binding sites is largely constitutive, MRTF is responsive to extracellular stimuli-elicited Rho-ROCK-actin signaling (Esnault et al., 2014; Gualdrini et al., 2016). To test actin dynamic-induced MRTF-SRF activity exerts direct transcriptional control of the molecular clock circuit, we determined MRTF or SRF chromatin occupancy on core clock genes. MRTF/SRF DNA binding element, CArG box, were identified within gene regulatory regions containing proximal promoter (+-2kb of transcription start site) of core clock genes using TRANSFAC. Using chromatin immunoprecipitation-qPCR (ChIP-qPCR), ∼8-10-fold of SRF enrichment on known CArG sites within *alpha-actin* and *vinculin* promoters were detected over IgG control, as expected (Fig. 8A). Similar degrees of ∼6-10-fold enrichment of SRF occupancy were found on clock gene regulatory regions within *Per1, Per2, Nr1d1*, and *Nfil3*, demonstrating these clock components as direct SRF target genes. We next performed immunoprecipitation with a MRTF-A antibody to further determine MRTF/SRF-mediated transcriptional control of these genes. As shown in Fig. 8B, we observed similar extent of MRTF-A chromatin association of the clock gene regulatory regions examined. Interestingly, MRTF-A occupancy of identified CArG site on Per2 promoter was markedly higher than that of SRF. To explore whether loss of SRF or MRTF affects their chromatin occupancy on clock target genes, we generated stable cell lines containing SRF or MRTF-A/B shRNA knockdown (Liu et al., 2020). Loss of *Srf* by stable knockdown in Per2-Luc fibroblasts largely abolished binding to identified regulatory regions of clock gene targets, with detected enrichment comparable to IgG control (Fig. 8C). Similar loss of MRTF-A binding to target promoters were observed in cells with stable expression of MRTF-A/B shRNA (Fig. 8D). Notably, the extent of MRTF-A enrichment was mostly attenuated in cells with scramble control expression as compared to normal Per2-Luc fibroblasts. Together, these findings identified direct MRTF/SRF transcriptional control within specific core clock components. Collectively, our results revealed a MRTF/SRF-mediated regulatory mechanism in controlling clock gene transcription, and this mechanism is responsive to extracellular niche signals transmitted *via* the Rho-ROCK-actin remodeling signaling cascade (Fig. 8E).

**Figure 8.**
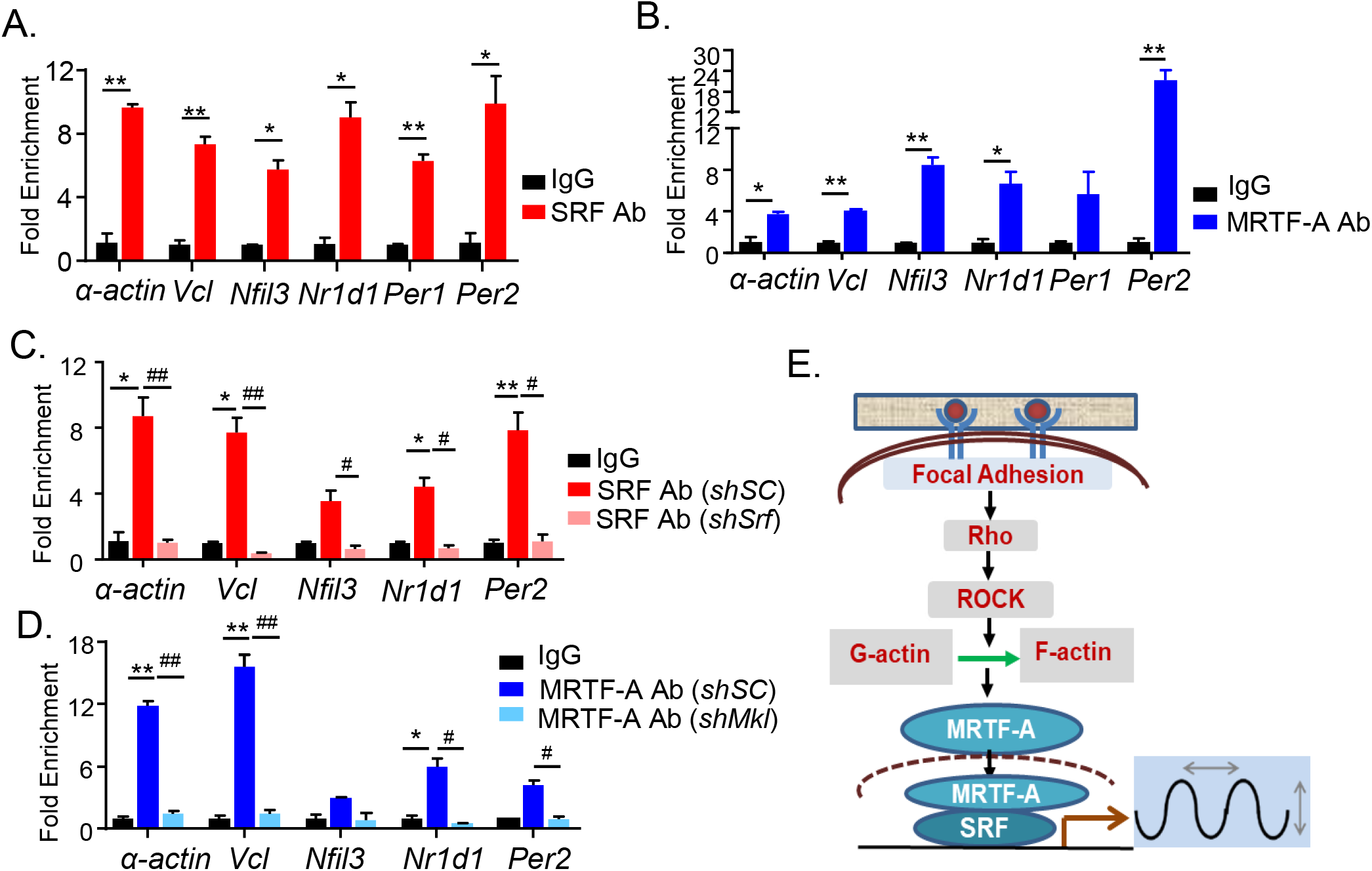
Direct MRTF/SRF transcriptional control of core clock genes. (A, B) ChIP-qPCR analysis of chromatin occupancy of SRF (A), and MRTF-A (B), on regulatory regions of core clock genes, as compared to IgG control. SRF and MRTF-A binding to known CArG sites of α-actin and vinculin (Vcl) promoters were as positive controls. (C, D) ChIP-qPCR analysis of SRF (C), and MRTF (D) chromatin occupancy in fibroblasts with stable expression of scrambled control (*shSC*), *Srf* shRNA knockdown (*shSrf*, C), or *Mrtf* shRNA knockdown (*shMrtf*, D). *, **: P≤0.05 or 0.01 SRF or MRTF-A antibody vs. IgG control. #, ##: P≤0.05 or 0.01 *shSrf* or *shMrtf* vs. *shSC*. (E) Schematic model of focal adhesion-actin cytoskeleton-MRTF/SRF signaling cascade in transducing cellular niche cues to circadian clock circuit.

## Discussion

Through systematic interrogation of distinct steps involved in integrin-actin cytoskeleton-MRTF/SRF signaling on circadian clock modulation, our current study established the role of this signal transduction cascade in linking circadian clock response to its extracellular physical niche. The actin cytoskeleton-MRTF/SRF pathway is a key mechanisms driving cellular developmental processes involving adhesion, migration, proliferation and differentiation (Gualdrini et al., 2016; Long et al., 2007; Olson and Nordheim, 2010). Our findings thus implicate circadian clock involvement in fundamental aspects of cellular behavior in tissue development, growth and remodeling processes.

Modulation of MRTF/SRF activity, by interfering with actin cytoskeleton organization and its up-stream signaling event, yielded largely consistent modulation of key clock properties. Chemicals disrupting actin polymerization, including Cyt D, Lat B and Cyt B, augmented cycling amplitude with longer period, whereas enhancing F-actin polymerization by Jasplakinolide led to opposite effects of reduced amplitude with shortened period. Blocking events up-stream of actin remodeling by inhibition of ROCK or FAK, or integrin blockade also led to increased period length and oscillation amplitude. Genetic inhibition of *Srf* or *Mrtf-a/b*, transcriptional transducers of ECM-actin dynamics-induced response, further corroborated these findings by demonstrating the direct effects of *Srf* and *Mrtf*. These interventions robustly influence clock gene transcription consistent with MRTF/SRF signaling activity, and we identified clock repressors, Per2, Cry2 and Nfil3 as direct transcriptional targets of SRF and MRTF.

Despite the mostly consistent modulatory effects observed among distinct interventions of the integrin-actin cytoskeleton-MRTF/SRF cascade on clock activity, the transcriptional response observed among specific clock components, including *CLOCK, Per1, Per2* and *Nr1d1*, differ. This could be due to the complexity of the interlocked transcriptional-translational feedback loop that drives the molecular clock circuitry, and the precise regulation of clock function by these interventions may depend on the distinct combined effects of transcriptional controls involved. Notably, the period lengthening and amplitude-enhancing effects of actin depolymerizing agents were comparable between established Per2::Luc reporter lines tested, mouse fibroblasts and U2OS osteosarcoma cell line. In contrast, Jasplakinolide induced a stronger response at low concentrations in U2OS than Per2::Luc mouse fibroblasts, potentially due to differing intrinsic sensitivity to actin polymerization between these cell types. The effect of blocking integrin-mediated adhesion signaling on clock, by RGD or FAK knockdown, are relatively moderate as compared to actin-disrupting molecules. As multitude of intracellular molecules and pathways are involved in transducing extracellular matrix cues to modulate actin dynamic-driven MRTF/SRF transactivation (Olson and Nordheim, 2010), it is conceivable that interfering with cell-matrix interactions through RGD or FAK alone may not be as robust as intervention of downstream actin remodeling or direct inhibition of MRTF/SRF activity.

As a time-keeping mechanism to anticipate and adapt to cyclic environmental stimuli, circadian clock are entrained to extrinsic and endogenous timing cues (Finger et al., 2020). Serum stimulation is a strong entrainment signal for peripheral clocks (Balsalobre et al., 1998). Gerber et al. demonstrated that serum-induced actin remodeling and modulation of MRTF/SRF activity mediates clock entrainment in liver (Gerber et al., 2013), and this pathway is responsive to upstream signaling events involving Rho kinase activation in fibroblasts (Esnault et al., 2014). Our new findings further extend these previous studies in elucidating how actin cytoskeleton-associated signaling transduce ECM-integrin-associated niche cues to MRTF/SRF-mediated transcriptional response that controls clock gene expression. Distinct integrins form linkages with intracellular actin cytoskeleton through interactions with specific extracellular matrix components (Maartens and Brown, 2015; Romero et al., 2020), thereby transducing ECM cues through actin-MRTF/SRF-mediated signaling (Posern and Treisman, 2006). Our findings linking integrin-associate signaling with clock modulation suggests that integrin may transduce specific physical niche cues from extracellular matrix to induce clock synchronization. Additionally, by culturing cells on matrices with defined physical properties, we demonstrated that softening of ECM led to progressive period lengthening with augmented cycling amplitude. Collectively, these findings suggest a potential circadian clock sensing mechanism of its immediate microenvironment involving integrin-mediated interactions with ECM and mechanical tension of matrix stiffness. These results are in line with recent reports from Yang et al. demonstrating that matrix stiffness modulates clock function in mammary epithelial cells (Yang et al., 2017), and that softer 3-D matrix induced stronger clock gene oscillations than stiff 2-D matrices (Williams et al., 2018). It is notable that in mammary epithelial cells, ROCK kinase inhibition by Y27632 was found to augment clock amplitude as we observed in both Per2-Luc fibroblasts and U2OS cells (Yang et al., 2017). Interestingly, fibroblasts from lung and mammary mesenchyme were reported to display inverse regulation by matrix rigidity (Williams et al., 2018), suggesting that physical properties of ECM, including stiffness or composition, may have distinct influence on clock functions in distinct cell types that may provide tissue-specific adaptations to unique extracellular environment (Streuli and Meng, 2019). Based on our findings, it will be intriguing to explore whether these cell type-dependent clock responses to matrix cues could be mediated by actin cytoskeleton-mediated MRTF/SRF transcriptional events, and further interrogated in diverse cellular or *in vivo* models.

Circadian clock modulates key stem cell behaviors in tissue remodeling processes involving epidermal, intestinal stem cells or myogenic progenitors (Chatterjee et al., 2013; Chatterjee et al., 2019; Chatterjee et al., 2014; Karpowicz et al., 2013; Plikus et al., 2013). Clock function is required for coordinating stem cell activation with environmental stimuli (Janich et al., 2011), and clock components, Bmal1 and Rev-erbα, exert antagonistic roles in muscle stem cell proliferation and differentiation during muscle regeneration (Chatterjee et al., 2013; Chatterjee et al., 2019; Chatterjee et al., 2014; Gao et al., 2020). Stem cell quiescence and self-renewal properties are intimately linked with its physical niche environment (McBeath et al., 2004). The mechanistic connection of clock modulation by its physical niche we uncovered implicates its potential contribution to stem cell properties in tissue development or remodeling known to involve ECM interactions (39). An interplay between cellular physical niche signals and the clock circuit may confer spatial and temporal coordination to direct stem cell behavior in developmental processes. Future studies are warranted to explore how an ECM-actin cytoskeleton-MRTF/SRF regulatory axis in modulating clock activity may apply to stem cell biology.

Our study revealed that small molecule modulators of actin dynamic and related signaling pathway significantly impact circadian clock function, mediated by MRTF/SRF transcriptional activity. It is conceivable, that agents such as RhoA/ROCK inhibitors or SRF modulators may have un-intended biological activities on clock modulation. On the other hand, these compounds or chemical derivatives may have potential applications in metabolic regulation or cancer therapy. With the recent effort in targeting clock for cancer and metabolic diseases (Chen et al., 2012; Cho et al., 2012; Dong et al., 2019; Oshima et al., 2019; Solt et al., 2012; Sulli et al., 2018), identification of the actin cytoskeleton-MRTF/SRF pathway in regulating circadian clock may provide a novel mechanistic basis for future studies.

## Materials and methods

### Cell Culture

Mouse primary fibroblasts originally obtained from enzymatic digestion of tail of mPER2::LUC-SV40 knock-in mice the mPer2 Luciferase-SV40 reporter (Welsh et al., 2004) that overcame replicative senescence. These cells and U2OS cell line containing Per2-Luciferase reporter were kind gifts from Dr. Steve Kay at University of Southern California. Cells were maintained in DMEM supplemented with 10% fetal bovine serum and antibiotics in a standard tissue culture incubator at 37^0^C with 5% CO2, as described previously (Liu et al., 2008). Cells were seeded in 24-well culture plates, grown to confluence, and switched to Lumicycle bioluminescence recording media for continuous Lumicycle recording for 7 days. Lumicycle recording media contains HEPES-buffered, air-equilibrated DMEM with 10 mM HEPES, 1.2 g/L NaHCO3, 25 U/ml penicillin, 25 g/ml streptomycin, 2% B-27, and 1 mM luciferin.

### Chemicals and Reagents

Cytochalasin D, Cytochalasin B, Latrunculin B, Jasplakinolide, and ROCK inhibitor Y27632 were purchased from Cayman Chemicals. Integrin RGD blocking peptide Cyclo [Arg-Gly-Asp-D-Phe-Val] and control Cyclo [Arg-Ala-Asp-D-Phe-Val] were obtained from Enzo Life Sciences. Electrospun scaffolds, poly(ε-caprolactone) (PCL) and polyether-ketoneketone (PEKK) were synthesized as described previously (Maldonado et al., 2015). The substrates were air plasma-treated and collagen-conjugated for cell adhesion, and mechanical property assessed by Young’s modulus was determined using atomic force microscopy. The scaffolds were sterilized and inserted in 24-well plates for cell culture.

### Luminometry, Bioluminescence Recording and Data Analysis

Measurement of bioluminescence rhythms from Per2-Luc mouse fibroblast cell culture were conducted using a luminometer LumiCycle 96 (Actimetrics), as described (Chen et al., 2012; Liu et al., 2008; Ramanathan et al., 2018). Briefly, 24-well tissue culture plates were sealed with plastic film and placed inside a bacterial incubator maintained at 36^0^C, 0% CO2. Luminescence from each well was measured for ∼70 s at intervals of 10 min and recorded as counts/second. 7 days of real-time bioluminescence recording data was analyzed using LumiCycle Analysis Program (Actimetrics) to determine clock oscillation period, length amplitude and phase. Briefly, raw data following the first cycle from day 2 to day 5 were fitted to a linear baseline, and the baseline-subtracted data (polynomial number = 1) were fitted to a sine wave, from which period length and goodness of fit and damping constant were determined. For samples that showed persistent rhythms, goodness-of-fit of >80% was usually achieved.

### siRNA and shRNA transfection, lentiviral plasmid construction and infection

siRNAs were purchased from Integrated DNA Technologies. Transfection were conducted using Lipofectamine RNAi MAX reagent (Invitrogen). 48-72 hours post transfection, cells were collected for protein or used for Lumicycle analysis. Lentiviral vectors expressing SRF or MRTF-A/B shRNA were obtained from Open Biosystems, as previously described (Liu et al., 2020). Three siRNA or shRNAs for each target were tested for knockdown efficiency. To generate shRNA knockdown lines, recombinant lentiviruses were produced by transient transfection in 293T cells using the calcium-phosphate method. Infectious lentiviral particles were harvested at 48 hr post-transfection to infect Per2-Luc fibroblasts. Two days post-infection, cells were selected with 2 μg/ml puromycin to obtain stable expression lines used for luminometry analyses. Lentiviral transduction efficiency was tested using lentiviral expressing GFP construct for GFP expression efficiency at nearly 95%.

### Immunoblot Analysis

20-40 µg of total protein were resolved on SDS-PAGE gel and transferred to PVDF membrane for immunoblotting (Liu et al., 2020). Immunoblots were developed by chemiluminescence kit (Pierce Biotechnology). Source and dilution information of primary antibodies are included as Suppl. Table S1. Appropriate specific secondary antibodies were used at a dilution of 1:3000.

### Quantitative real-time PCR analysis

RNeasy miniprep kits (Qiagen) were used to isolate total RNA from cells. cDNA was generated using q-Script cDNA kit (Quanta Biosciences) and quantitative PCR was performed on an ABI Light Cycler with SYBR Green (Quanta Biosciences). Relative mRNA expression was determined using the comparative Ct method to normalize target genes 36B4 as internal controls. Primers are designed using Primer Bank experimentally validated sequences. Primer sequences are listed in supplementary Table S2.

### Phalloidin staining of F-actin

Fluorescence staining of actin was performed using Alexa Fluor 488-conjugateed phalloidin, similarly as described (Liu et al., 2020). Briefly, cells were fixed with 3.7% formaldehyde and permeabilized using 0.1% Triton X-100. Alexa Fluor 488 phalloidin (10 µg/mL) at 1:200 dilution together with DAPI (1:2000) was incubated for 30 min at room temperature. Fluorescence images were taken using an ECHO microscope.

### ChIP-qPCR

Immunoprecipitation was performed using SRF or MKL1 antibody or control rabbit IgG with Magnetic Protein A/G beads (Magna ChIP A/G kit, Millipore), as described previously (Chatterjee et al., 2013). Mouse myofibroblast chromatin was sonicated and purified following formaldehyde fixation. Real-time PCR was carried out in triplicates using purified chromatin with specific primers for predicted Bmal1 binding E or E’-box elements identified within the gene regulatory regions. Primers flanking known Bmal1 E-box in Rev-erbα promoter was used as positive control and TBP first exon primers as negative control. Fold enrichment were expressed normalized to 1% of input over IgG. ChIP primer sequences are listed in Supplementary Table S3.

### Statistical analysis

Data was expressed as mean ± SE. Differences between groups were examined for statistical significance using unpaired two-tailed Student’s t-test or ANOVA for multiple group comparison as indicated using Prism by GraphPad. P<0.05 was considered statistically significant.

## Acknowledgements

We thank Drs. Steve Kay and Meng Qu at the University of Southern California for sharing the luciferase reporter cell lines used in this study. KM is a faculty member supported by the NCI-designated Comprehensive Cancer Center at the City of Hope National Cancer Center. This project was supported by a grant from National Institute of Health 1R01DK112794 to KM. The funder had no role in study design, data collection and analysis, decision to publish, or preparation of the manuscript.

## Authorship Statement

XX and WL: data curation and investigation, formal analysis, manuscript editing; JN, MQ and SK: data curation and manuscript editing; KM: formal analysis, project administration, manuscript writing and funding acquisition.

## Competing Interests

The authors declare that no competing interests exist that is relevant to the subject matter or materials included in this work.

## Figure Legends

**Fig. S1.**
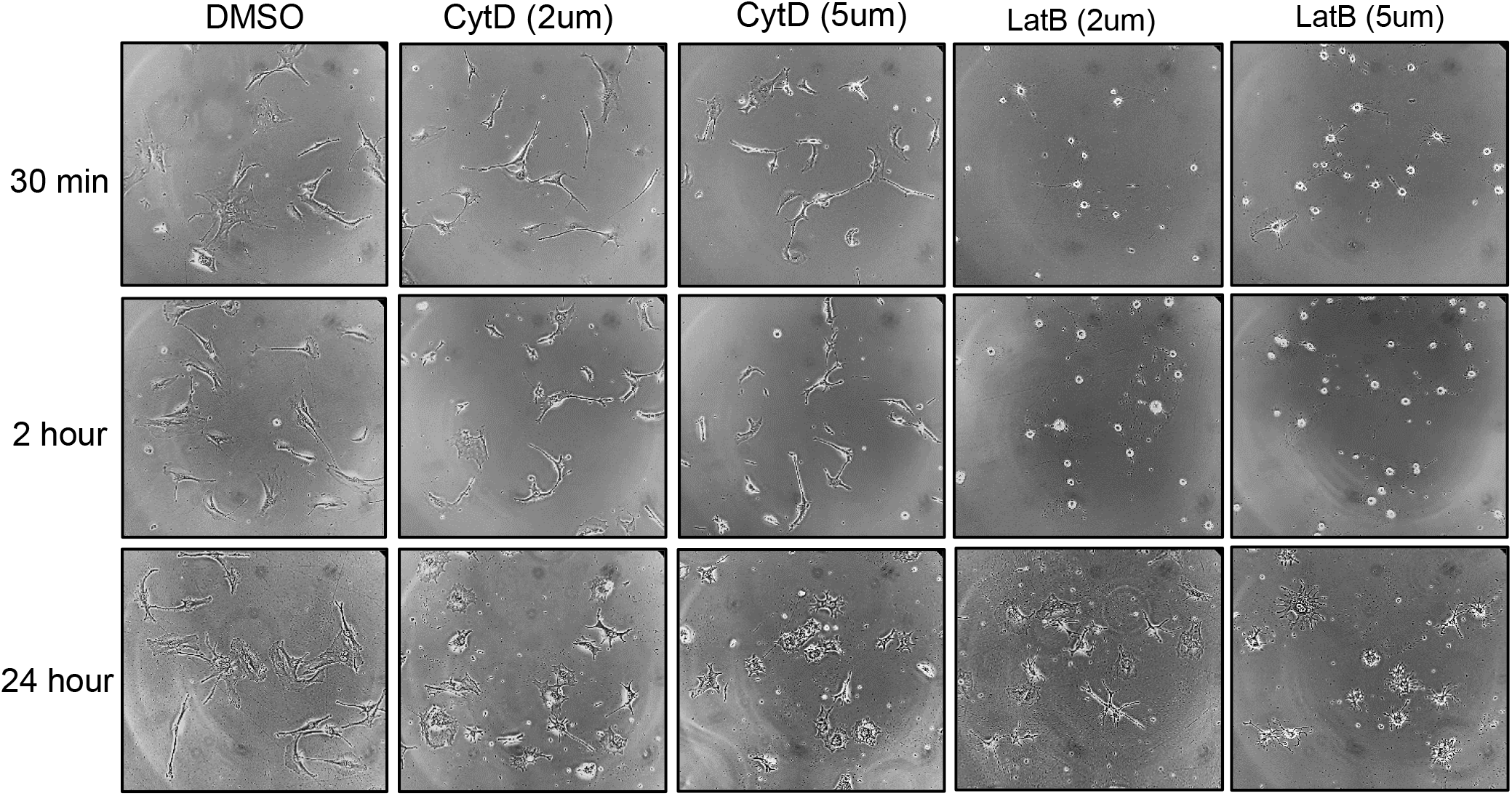
Representative phase contrast images of cell morphology in response to Cytochalasin D (Cyt D) and Latrunculin B (Lat B) treatment at indicated concentrations and time course.

**Fig. S2.**
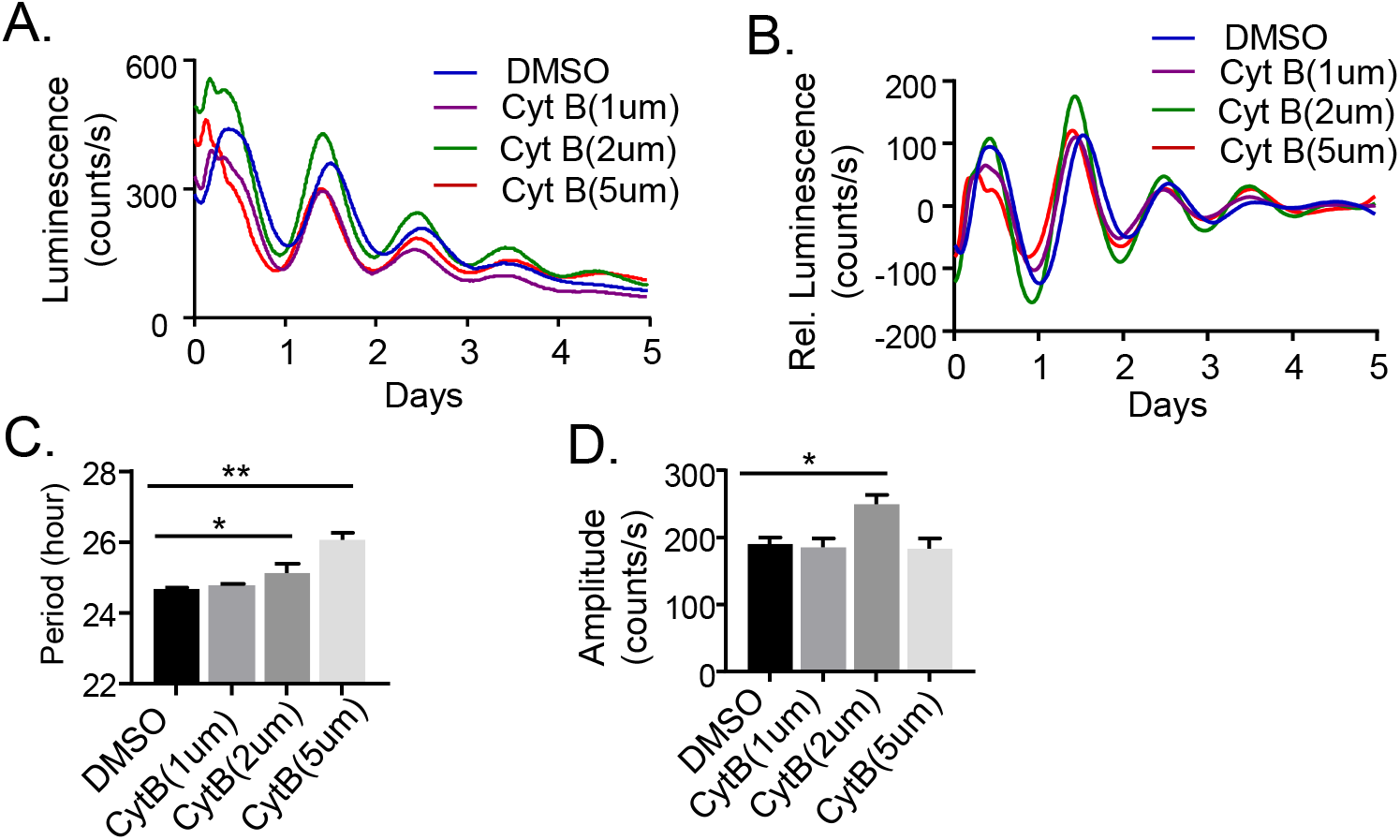
Effect of Cytochalasin B on clock modulation. (A, B). The average original bioluminescence plots (A), and baseline-subtracted plots (B) of Per2-Luciferase knock-in mouse fibroblasts subjected to Cytochalasin B (CytB) treatment at indicated concentrations, with quantitative analysis of clock period length (C) and amplitude (D). N=3/treatment group.

**Fig. S3.**
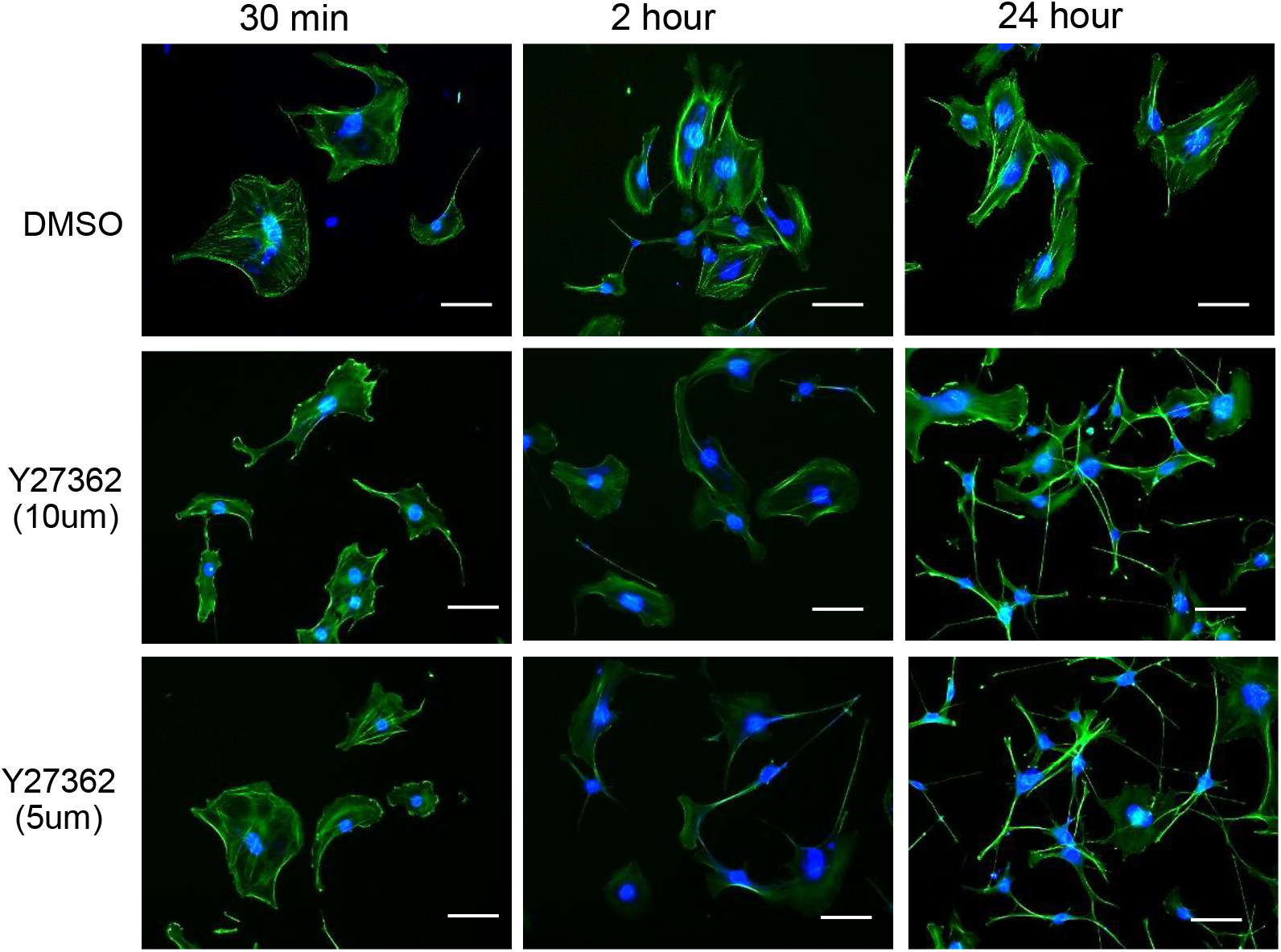
Representative fluorescence images of phalloidin staining of actin stress fibers in control (DMSO) or Y27362-treated Per-2 knock-in fibroblasts at indicated concentrations and time course. Scale bar: 100 μm.

**Fig. S4.**
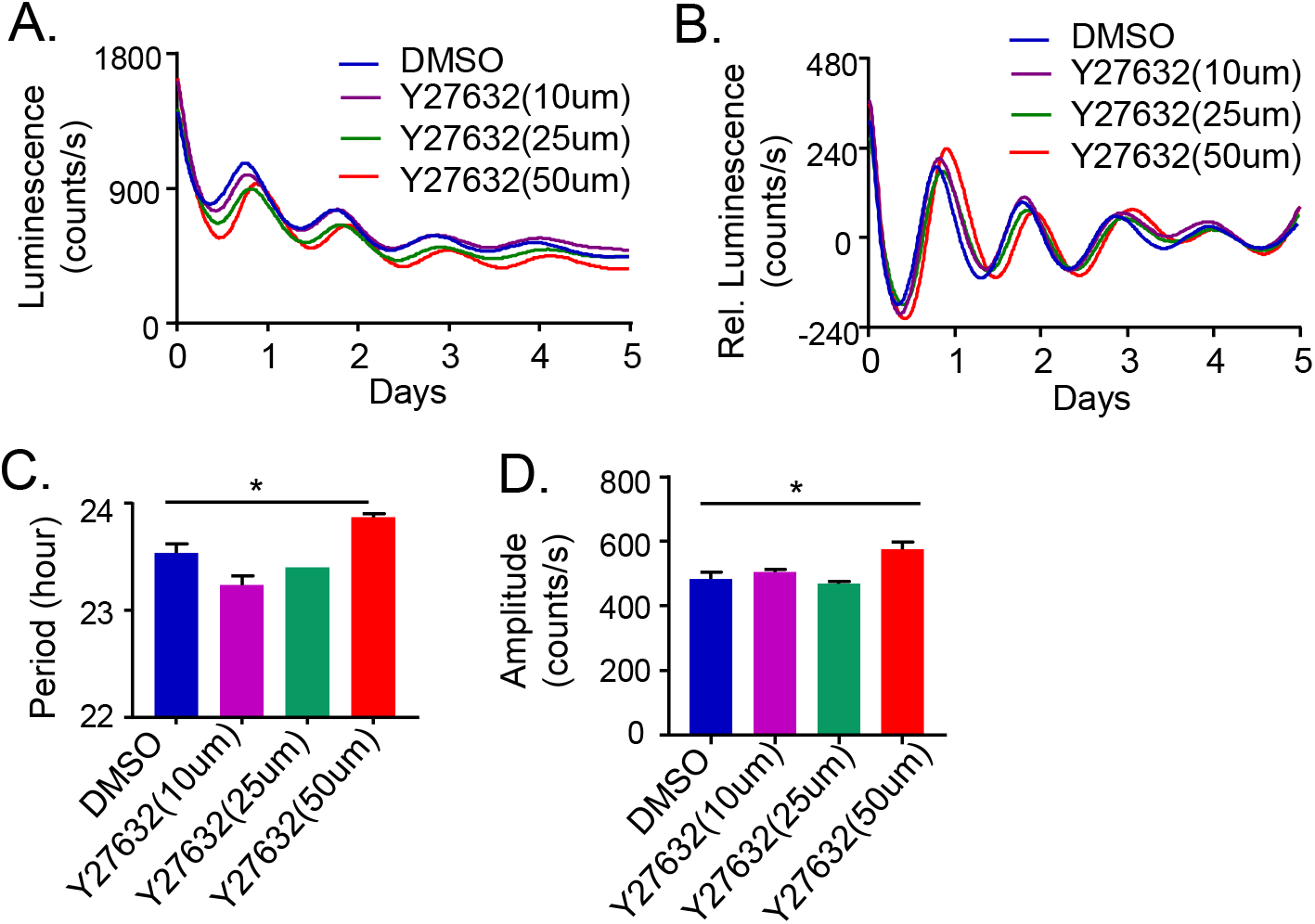
Effect of ROCK inhibitor Y27632 on clock modulation in Per2::Luc U2OS cells. The average original plots of bioluminescence (A), and baseline-subtracted plots (B) of Per2-Luciferase knock-in U2OS cells subjected to Y27632 treatment at indicated concentrations, with quantitative analysis of clock period length (C) and amplitude (D). N=3/treatment group.

**Supplemental Table 1.** Primary antibodies list.

| Antibody | Source | Cat# | Dilution |
| --- | --- | --- | --- |
| SRF | Santa Cruz | SC-25290 | 1:1000 |
| MRTF-A (MKL1) | Santa Cruz | SC-390324 | 1:1000 |
| FAK | Santa Cruz | SC-557 | 1:1000 |
| GAPDH | ThermoFisher | AM4300 | 1:3000 |

**Supplemental Table 2.**
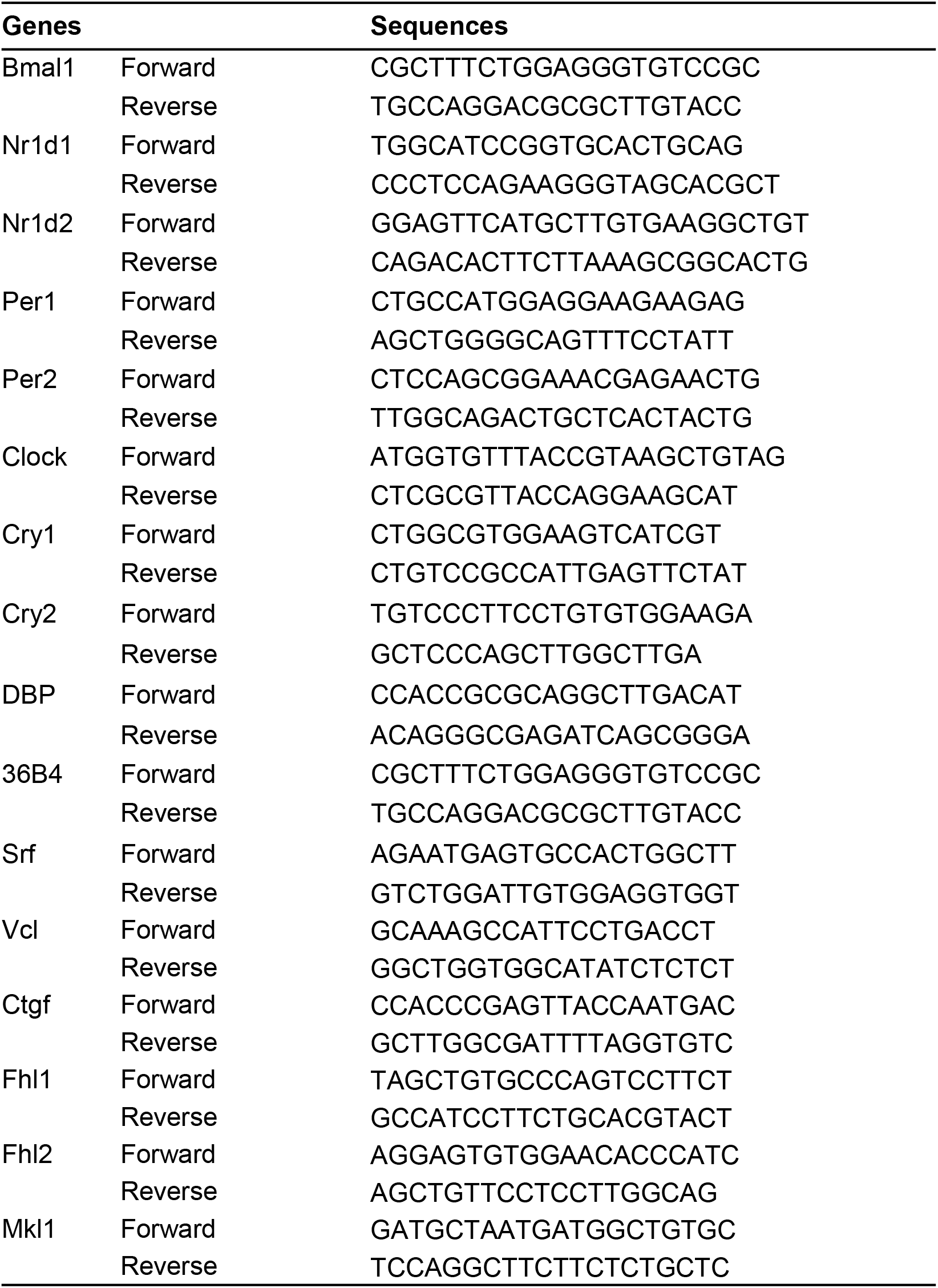
Primer sequence for qPCR analysis.

**Supplemental Table 3.**
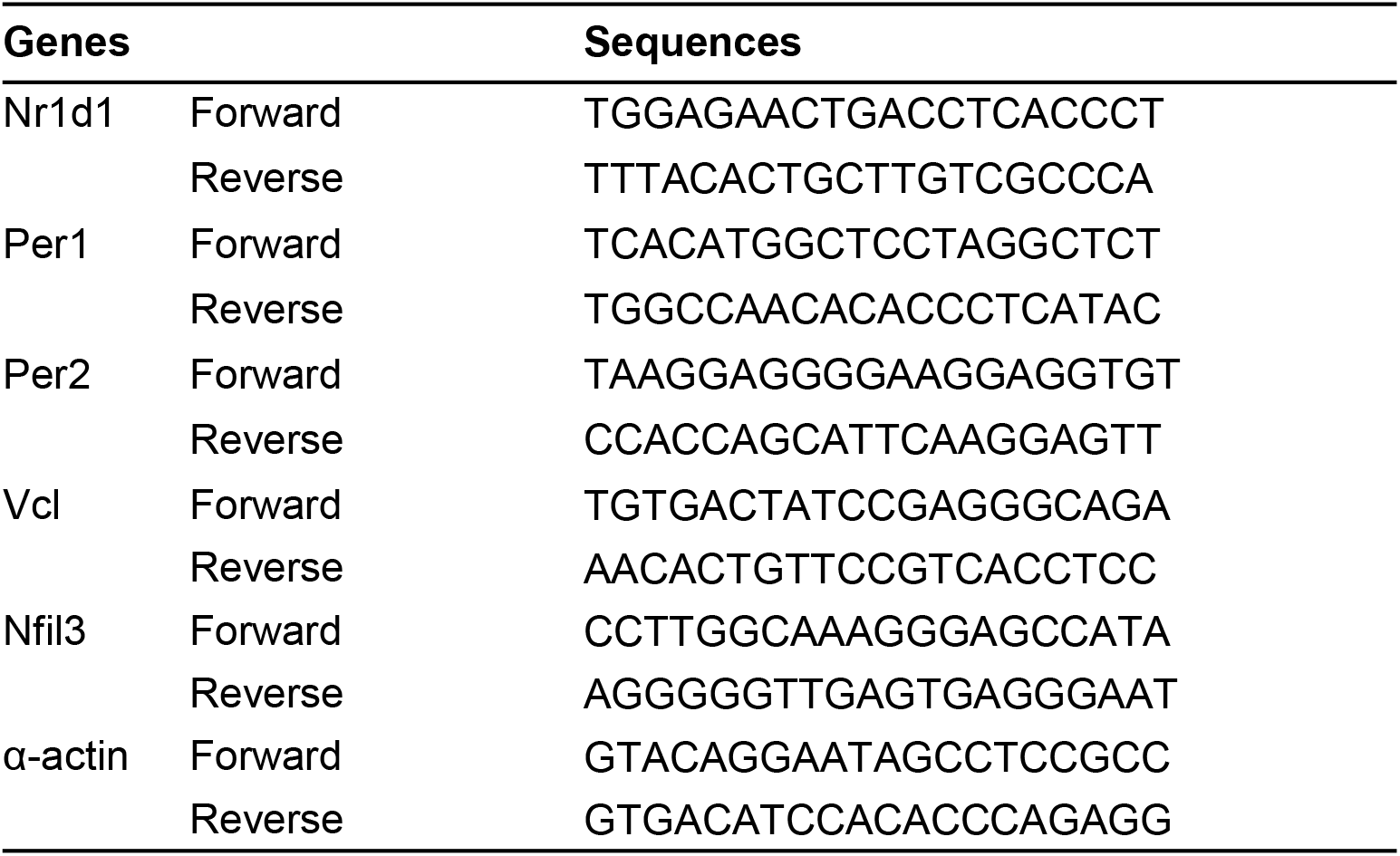
Primer sequence for ChIP-qPCR analysis.

## Notes

### Competing Interest Statement

The authors have declared no competing interest.

## Reference

Allingham, J. S., Klenchin, V. A. and Rayment, I. (2006). Actin-targeting natural products: structures, properties and mechanisms of action. Cell Mol Life Sci 63, 2119–34.

Balsalobre, A., Damiola, F. and Schibler, U. (1998). A serum shock induces circadian gene expression in mammalian tissue culture cells. Cell 93, 929–37.

Buxton, O. M., Cain, S. W., O’Connor, S. P., Porter, J. H., Duffy, J. F., Wang, W., Czeisler, C. A. and Shea, S. A. (2012). Adverse metabolic consequences in humans of prolonged sleep restriction combined with circadian disruption. Sci Transl Med 4, 129ra43.

Chatterjee, S., Nam, D., Guo, B., Kim, J. M., Winnier, G. E., Lee, J., Berdeaux, R., Yechoor, V. K. and Ma, K. (2013). Brain and muscle Arnt-like 1 is a key regulator of myogenesis. J Cell Sci 126, 2213–24.

Chatterjee, S., Yin, H., Li, W., Lee, J., Yechoor, V. K. and Ma, K. (2019). The Nuclear Receptor and Clock Repressor Rev-erbalpha Suppresses Myogenesis. Sci Rep 9, 4585.

Chatterjee, S., Yin, H., Nam, D., Li, Y. and Ma, K. (2014). Brain and muscle Arnt-like 1 promotes skeletal muscle regeneration through satellite cell expansion. Exp Cell Res.

Chen, Z., Yoo, S. H., Park, Y. S., Kim, K. H., Wei, S., Buhr, E., Ye, Z. Y., Pan, H. L. and Takahashi, J. S. (2012). Identification of diverse modulators of central and peripheral circadian clocks by high-throughput chemical screening. Proc Natl Acad Sci U S A 109, 101–6.

Cho, H., Zhao, X., Hatori, M., Yu, R. T., Barish, G. D., Lam, M. T., Chong, L. W., DiTacchio, L., Atkins, A. R., Glass, C. K. et al. (2012). Regulation of circadian behaviour and metabolism by REV-ERB-alpha and REV-ERB-beta. Nature 485, 123–7.

Dibner, C., Schibler, U. and Albrecht, U. (2010). The mammalian circadian timing system: organization and coordination of central and peripheral clocks. Annu Rev Physiol 72, 517–49.

Dong, Z., Zhang, G., Qu, M., Gimple, R. C., Wu, Q., Qiu, Z., Prager, B. C., Wang, X., Kim, L. J. Y., Morton, A. R. et al. (2019). Targeting Glioblastoma Stem Cells through Disruption of the Circadian Clock. Cancer Discov 9, 1556–1573.

Esnault, C., Stewart, A., Gualdrini, F., East, P., Horswell, S., Matthews, N. and Treisman, R. (2014). Rho-actin signaling to the MRTF coactivators dominates the immediate transcriptional response to serum in fibroblasts. Genes Dev 28, 943–58.

Finger, A. M., Dibner, C. and Kramer, A. (2020). Coupled network of the circadian clocks: a driving force of rhythmic physiology. FEBS Lett 594, 2734–2769.

Gao, H., Xiong, X., Lin, Y., Chatterjee, S. and Ma, K. (2020). The clock regulator Bmal1 protects against muscular dystrophy. Exp Cell Res 397, 112348.

Gerber, A., Esnault, C., Aubert, G., Treisman, R., Pralong, F. and Schibler, U. (2013). Blood-borne circadian signal stimulates daily oscillations in actin dynamics and SRF activity. Cell 152, 492–503.

Gualdrini, F., Esnault, C., Horswell, S., Stewart, A., Matthews, N. and Treisman, R. (2016). SRF Co-factors Control the Balance between Cell Proliferation and Contractility. Mol Cell 64, 1048–1061.

Hill, C. S., Wynne, J. and Treisman, R. (1995). The Rho family GTPases RhoA, Rac1, and CDC42Hs regulate transcriptional activation by SRF. Cell 81, 1159–70.

Janich, P., Pascual, G., Merlos-Suarez, A., Batlle, E., Ripperger, J., Albrecht, U., Cheng, H. Y., Obrietan, K., Di Croce, L. and Benitah, S. A. (2011). The circadian molecular clock creates epidermal stem cell heterogeneity. Nature 480, 209–14.

Karatsoreos, I. N., Bhagat, S., Bloss, E. B., Morrison, J. H. and McEwen, B. S. (2011). Disruption of circadian clocks has ramifications for metabolism, brain, and behavior. Proc Natl Acad Sci U S A 108, 1657–62.

Karpowicz, P., Zhang, Y., Hogenesch, J. B., Emery, P. and Perrimon, N. (2013). The circadian clock gates the intestinal stem cell regenerative state. Cell Rep 3, 996–1004.

Kettner, N. M., Katchy, C. A. and Fu, L. (2014). Circadian gene variants in cancer. Ann Med 46, 208–20.

Lin, H. H. and Farkas, M. E. (2018). Altered Circadian Rhythms and Breast Cancer: From the Human to the Molecular Level. Front Endocrinol (Lausanne) 9, 219.

Liu, A. C., Tran, H. G., Zhang, E. E., Priest, A. A., Welsh, D. K. and Kay, S. A. (2008). Redundant function of REV-ERBalpha and beta and non-essential role for Bmal1 cycling in transcriptional regulation of intracellular circadian rhythms. PLoS Genet 4, e1000023.

Liu, R., Xiong, X., Nam, D., Yechoor, V. and Ma, K. (2020). SRF-MRTF signaling suppresses brown adipocyte development by modulating TGF-beta/BMP pathway. Mol Cell Endocrinol 515, 110920.

Long, X., Creemers, E. E., Wang, D. Z., Olson, E. N. and Miano, J. M. (2007). Myocardin is a bifunctional switch for smooth versus skeletal muscle differentiation. Proc Natl Acad Sci U S A 104, 16570–5.

Maartens, A. P. and Brown, N. H. (2015). Anchors and signals: the diverse roles of integrins in development. Curr Top Dev Biol 112, 233–72.

Maldonado, M., Ico, G., Low, K., Luu, R. J. and Nam, J. (2016). Enhanced Lineage-Specific Differentiation Efficiency of Human Induced Pluripotent Stem Cells by Engineering Colony Dimensionality Using Electrospun Scaffolds. Adv Healthc Mater 5, 1408–12.

Maldonado, M., Luu, R. J., Ico, G., Ospina, A., Myung, D., Shih, H. P. and Nam, J. (2017). Lineage-and developmental stage-specific mechanomodulation of induced pluripotent stem cell differentiation. Stem Cell Res Ther 8, 216.

Maldonado, M., Wong, L. Y., Echeverria, C., Ico, G., Low, K., Fujimoto, T., Johnson, J. K. and Nam, J. (2015). The effects of electrospun substrate-mediated cell colony morphology on the self-renewal of human induced pluripotent stem cells. Biomaterials 50, 10–9.

McBeath, R., Pirone, D. M., Nelson, C. M., Bhadriraju, K. and Chen, C. S. (2004). Cell shape, cytoskeletal tension, and RhoA regulate stem cell lineage commitment. Dev Cell 6, 483–95.

Miano, J. M. (2003). Serum response factor: toggling between disparate programs of gene expression. J Mol Cell Cardiol 35, 577–93.

Mocellin, S., Tropea, S., Benna, C. and Rossi, C. R. (2018). Circadian pathway genetic variation and cancer risk: evidence from genome-wide association studies. BMC Med 16, 20.

Olson, E. N. and Nordheim, A. (2010). Linking actin dynamics and gene transcription to drive cellular motile functions. Nat Rev Mol Cell Biol 11, 353–65.

Oshima, T., Niwa, Y., Kuwata, K., Srivastava, A., Hyoda, T., Tsuchiya, Y., Kumagai, M., Tsuyuguchi, M., Tamaru, T., Sugiyama, A. et al. (2019). Cell-based screen identifies a new potent and highly selective CK2 inhibitor for modulation of circadian rhythms and cancer cell growth. Sci Adv 5, eaau9060.

Plikus, M. V., Vollmers, C., de la Cruz, D., Chaix, A., Ramos, R., Panda, S. and Chuong, C.M. (2013). Local circadian clock gates cell cycle progression of transient amplifying cells during regenerative hair cycling. Proc Natl Acad Sci U S A 110, E2106–15.

Posern, G. and Treisman, R. (2006). Actin’ together: serum response factor, its cofactors and the link to signal transduction. Trends Cell Biol 16, 588–96.

Preitner, N., Damiola, F., Lopez-Molina, L., Zakany, J., Duboule, D., Albrecht, U. and Schibler, U. (2002). The orphan nuclear receptor REV-ERBalpha controls circadian transcription within the positive limb of the mammalian circadian oscillator. Cell 110, 251–60.

Ramanathan, C., Kathale, N. D., Liu, D., Lee, C., Freeman, D. A., Hogenesch, J. B., Cao, R. and Liu, A. C. (2018). mTOR signaling regulates central and peripheral circadian clock function. PLoS Genet 14, e1007369.

Romero, S., Le Clainche, C. and Gautreau, A. M. (2020). Actin polymerization downstream of integrins: signaling pathways and mechanotransduction. Biochem J 477, 1–21.

Scheer, F. A., Hilton, M. F., Mantzoros, C. S. and Shea, S. A. (2009). Adverse metabolic and cardiovascular consequences of circadian misalignment. Proc Natl Acad Sci U S A 106, 4453–8.

Schibler, U. and Sassone-Corsi, P. (2002). A web of circadian pacemakers. Cell 111, 919–22.

Shi, S. Q., Ansari, T. S., McGuinness, O. P., Wasserman, D. H. and Johnson, C. H. (2013). Circadian disruption leads to insulin resistance and obesity. Curr Biol 23, 372–81.

Solt, L. A., Wang, Y., Banerjee, S., Hughes, T., Kojetin, D. J., Lundasen, T., Shin, Y., Liu, J., Cameron, M. D., Noel, R. et al. (2012). Regulation of circadian behaviour and metabolism by synthetic REV-ERB agonists. Nature 485, 62–8.

Streuli, C. H. and Meng, Q. J. (2019). Influence of the extracellular matrix on cell-intrinsic circadian clocks. J Cell Sci 132.

Sulli, G., Rommel, A., Wang, X., Kolar, M. J., Puca, F., Saghatelian, A., Plikus, M. V., Verma, I. M. and Panda, S. (2018). Pharmacological activation of REV-ERBs is lethal in cancer and oncogene-induced senescence. Nature 553, 351–355.

Sun, Q., Chen, G., Streb, J. W., Long, X., Yang, Y., Stoeckert, C. J., Jr. and Miano, J. M. (2006). Defining the mammalian CArGome. Genome Res 16, 197–207.

Takahashi, J. S. (2017). Transcriptional architecture of the mammalian circadian clock. Nat Rev Genet 18, 164–179.

Turek, F. W., Joshu, C., Kohsaka, A., Lin, E., Ivanova, G., McDearmon, E., Laposky, A., Losee-Olson, S., Easton, A., Jensen, D. R. et al. (2005). Obesity and metabolic syndrome in circadian Clock mutant mice. Science 308, 1043–5.

Wang, D. Z., Li, S., Hockemeyer, D., Sutherland, L., Wang, Z., Schratt, G., Richardson, J. A., Nordheim, A. and Olson, E. N. (2002). Potentiation of serum response factor activity by a family of myocardin-related transcription factors. Proc Natl Acad Sci U S A 99, 14855–60.

Wei, L., Wang, L., Carson, J. A., Agan, J. E., Imanaka-Yoshida, K. and Schwartz, R. J. (2001). beta1 integrin and organized actin filaments facilitate cardiomyocyte-specific RhoA-dependent activation of the skeletal alpha-actin promoter. FASEB J 15, 785–96.

Welsh, D. K., Yoo, S. H., Liu, A. C., Takahashi, J. S. and Kay, S. A. (2004). Bioluminescence imaging of individual fibroblasts reveals persistent, independently phased circadian rhythms of clock gene expression. Curr Biol 14, 2289–95.

Williams, J., Yang, N., Wood, A., Zindy, E., Meng, Q. J. and Streuli, C. H. (2018). Epithelial and stromal circadian clocks are inversely regulated by their mechano-matrix environment. J Cell Sci 131.

Yang, N., Williams, J., Pekovic-Vaughan, V., Wang, P., Olabi, S., McConnell, J., Gossan, N., Hughes, A., Cheung, J., Streuli, C. H. et al. (2017). Cellular mechano-environment regulates the mammary circadian clock. Nat Commun 8, 14287.

Yoo, S. H., Yamazaki, S., Lowrey, P. L., Shimomura, K., Ko, C. H., Buhr, E. D., Siepka, S. M., Hong, H. K., Oh, W. J., Yoo, O. J. et al. (2004). PERIOD2::LUCIFERASE real-time reporting of circadian dynamics reveals persistent circadian oscillations in mouse peripheral tissues. Proc Natl Acad Sci U S A 101, 5339–46.

